# A complex structure of arrestin-2 bound to a G protein-coupled receptor

**DOI:** 10.1101/822957

**Authors:** Wanchao Yin, Zhihai Li, Mingliang Jin, Yu-Ling Yin, Parker W. de Waal, Kuntal Pal, Yanting Yin, Xiang Gao, Yuanzheng He, Jing Gao, Xiaoxi Wang, Yan Zhang, Hu Zhou, Karsten Melcher, Yi Jiang, Yao Cong, X. Edward Zhou, Xuekui Yu, H. Eric Xu

## Abstract

Arrestins comprise a family of signal regulators of G-protein-coupled receptors (GPCRs), which include arrestins 1 to 4. While arrestins 1 and 4 are visual arrestins dedicated to rhodopsin, arrestins 2 and 3 (Arr2 and Arr3) are β-arrestins known to regulate many nonvisual GPCRs. The dynamic and promiscuous coupling of Arr2 to nonvisual GPCRs has posed technical challenges to tackle the basis of arrestin binding to GPCRs. Here we report the structure of Arr2 in complex with neurotensin receptor 1 (NTSR1), which reveals an overall assembly that is strikingly different from the visual arrestin-rhodopsin complex by a 90° rotation of Arr2 relative to the receptor. In this new configuration, intracellular loop 3 (ICL3) and transmembrane helix 6 (TM6) of the receptor are oriented toward the N-terminal domain of the arrestin, making it possible for GPCRs that lack the C-terminal tail to couple Arr2 through their ICL3. Molecular dynamics simulation and crosslinking data further support the assembly of the Arr2–NTSR1 complex. Sequence analysis and homology modeling suggest that the Arr2–NTSR1 complex structure may provide an alternative template for modeling arrestin-GPCR interactions.

## Introduction

G-protein-coupled receptors (GPCRs) comprise the largest family of cell surface receptors that are involved in many aspects of human physiology and account for one-third of drug targets^1–3^. The functions of GPCRs are primarily mediated by two major families of signal transducers, G proteins and arrestins. The arrestin family contains four members, including two visual arrestins, arrestin-1(Arr1) and arestin-4(Arr4), and two β-arrestins, arrestin-2 (Arr2) and arrestin-3 (Arr3), also known as β-arrestin-1 and β-arestin-2, respectively. While visual arrestins are exclusively involved in signal transduction in vision through binding to rhodopsin, Arr2 and Arr3 are required for regulating signal transduction of many nonvisual GPCRs^4,5^. Binding of Arr2 and Arr3 to GPCRs not only blocks G protein binding but also mediates receptor endocytosis and numerous G protein-independent signaling pathways. Ligands that can selectively modulate arrestin-specific pathways have become a new paradigm of drug discovery as they could deliver more effective and safer treatments for a wide variety of diseases. As such, arrestin binding to GPCRs and its biased signaling have become a focus area of GPCR research^6^.

The mechanism of visual arrestin binding to rhodopsin has been illustrated by a femtosecond X-ray laser structure of the human rhodopsin bound to Arr1^7–10^. Arr1 has a crescent shaped conformation with a pseudo-twofold symmetry between the N- and C-domains of arrestin but is engaged with rhodopsin in an asymmetric fashion, with its finger loop inserted into the core cavity of the rhodopsin transmembrane domain (TMD). The complex is further stabilized by the binding of the phosphorylated C-terminal tail of rhodopsin to the positively charged N-terminal domain of arrestin. In this asymmetric arrangement, the C-edge loops of visual arrestin are positioned toward the membrane layer to form direct interactions with lipids. However, the situation of β-arrestin binding to nonvisual GPCRs is much more complex than that of visual arrestin binding to rhodopsin due to dynamic and promiscuous coupling of β-arrestin to many more GPCRs. Earlier studies suggested that β-arrestin can bind to GPCRs in multiple conformational states, including a core engagement state that is similar to the Arr1-rhodopsin complex, and a tail-engagement state whose interaction with arrestin is exclusively mediated by the receptor C-terminal tail^11^. Because of the promiscuous binding and highly dynamic complex assembly, the structure determination of an arrestin complex with a nonvisual GPCR is technically challenging and is a long-sought goal in the field of GPCR structural biology.

Neurotensin receptor 1 (NTSR1) belongs to the class A GPCR subfamily. When stimulated by a neurotensin peptide, NTSR1 couples to a G protein or arrestin for signal transduction that is essential for neurotransmission and neuromodulation in the central nervous system^12,13^ (Fig. 1a). It has been proposed as an important therapeutic target for a range of diseases including hypotension, obesity, analgesia, drug abuse, cancer, Parkinson’s disease and schizophrenia. Several structures of NTSR1 in complex with peptide agonists and inhibitory G protein have been reported, providing a detailed mechanism of ligand recognition and G protein coupling^14–16^. In this paper, we report a complex structure of NTSR1 bound to Arr2 solved by single particle cryo-electron microscopy (EM). This structure reveals an overall assembly of the Arr2-GPCR complex that is distinct from the Arr1-rhodopsin complex. Superposition of the two complex structures shows that the arrestins are in two orientations, which are 90° different around an axis vertical to the membrane layer. In this configuration, TM5/6 and ICL3 of NTSR1 are located close to the N-terminal region of Arr2, which allows ICL3 to interact with the N-domain of Arr2. Sequence alignment and homology modeling suggest that the Arr2–NTSR1 complex structure may serve as an alternative model for understanding arrestin interaction with nonvisual GPCRs.

**Figure 1.**
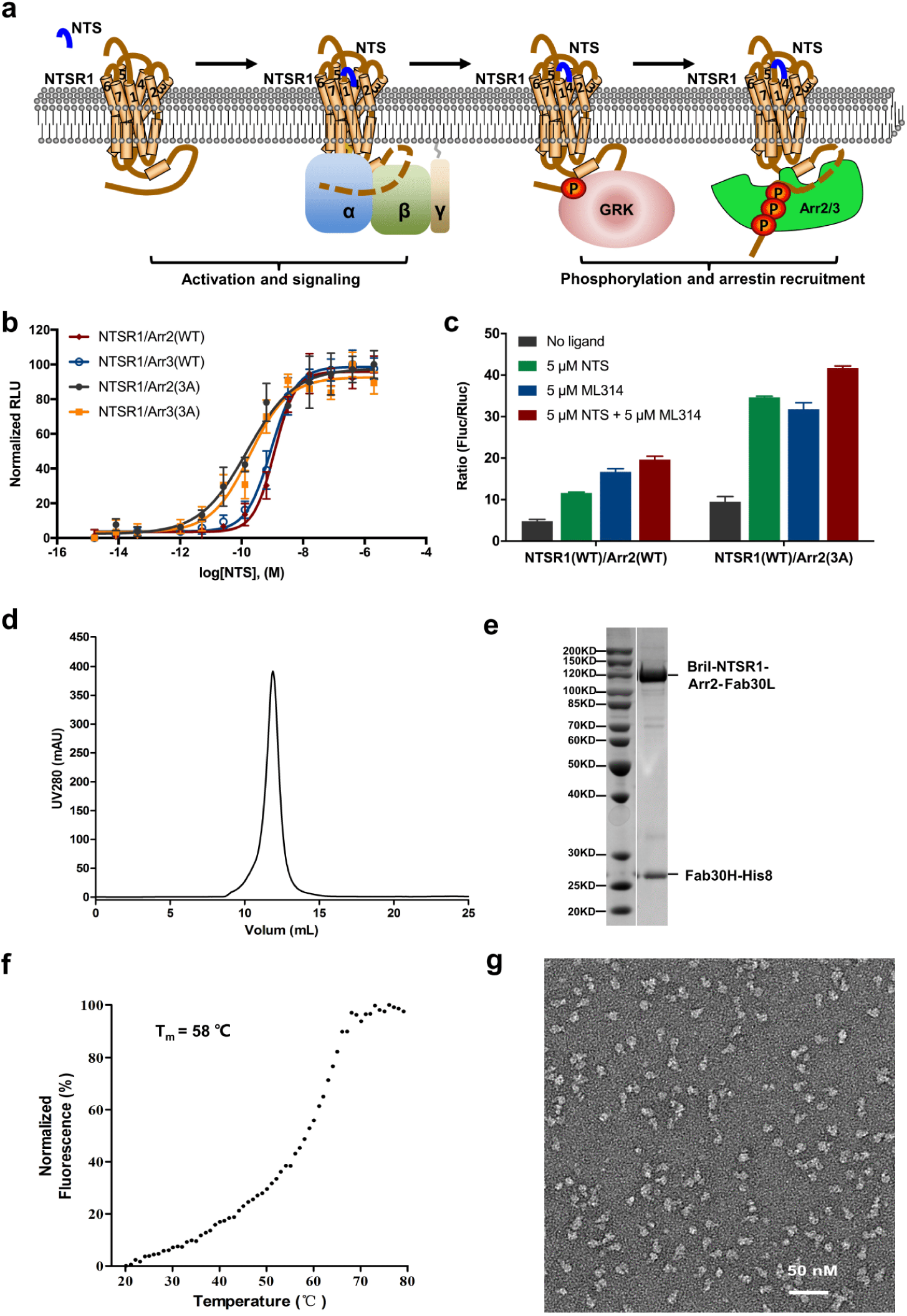
Characterization and structure determination of β-arrestin-NTSR1 complex a. Cartoon presentation of NTSR1 signaling mediated by G protein and β-arrestin b. β-arrestin-NTSR1 interaction promoted by addition of NTS (3-13) determined by NanoBiT assay. Symbols and error bars represent mean and s.e.m. of indicated independent numbers of experiments performed in triplicate. c. Neurotensin and ML314 improving Arr2–NTSR1 interaction determined by Tango assay d. Gel filtration profile of BRIL–NTSR1–Arr2–Fab30 complex e. SDS gel of the purified complex protein f. Thermal stability shift analysis of BRIL–NTSR1–Arr2–Fab30 fusion complex indicated a high thermostability of this complex with its melting temperature at 58°C g. Negative staining electron microscopy of BRIL–NTSR1–Arr2–Fab30 fusion complex.

### Functional characterization and structure determination of Arr2–NTSR1 complex

Interactions of arrestins to nonvisual GPCR is relatively weak and highly dynamic but obtaining a stable complex is an essential step for structural studies. Therefore, prior to our structural studies, we characterized the interaction of NTSR1 with arrestins by utilizing the commercial NanoBiT system, which is an effective method to detect protein-protein interactions^17^. NTSR1 was found to interact with both Arr2 and Arr3, which is enhanced by the increased concentrations of peptide agonist, neurotensin (NTS). Arrestin mutants with three C-terminal hydrophobic residues mutated to alanine (3A mutants; I386A, V387A and F388A in Arr2, and I387A, V388A and F389A in Arr3)^18,19^, which are known as pre-activated form of arrestins, showed increased binding activity to NTSR1 (Fig 1b).

We further tested arrestin-NTSR1 interaction by Tango assay, a cell-based assay that determines binding activities of membrane proteins to soluble binding partners^7,20^. Our Tango assays showed that the binding activity of NTSR1 to Arr2 was increased by about two to three fold, respectively, when peptide agonist NTS or small molecule agonist ML314, a known positive-allosteric modulator (PAM)^21,22^, was added. The binding activity of Arr2 to NTSR1 was increased by about four-fold when both NTS and ML314 were used. The binding of NTSR1 to Arr2 was further enhanced by introducing 3A mutations (Fig. 1c).

Based on above results, we performed structural studies using the 3A mutant of Arr2 in complex with NTSR1 in the presence of NTS and ML314 but the complex dissociated in cryo-EM grids due to the adverse effects associated with sample vitrification. To overcome this dissociation problem, we fused the wild type human NTSR1 with the human 3A mutant Arr2 at its C-terminus with a three amino acid linker (GSA). Cytochrome b562 RIL domain (BRIL) was also fused to the N-terminus of the receptor to increase the complex expression. Arr2 was further stabilized by fusing Fab30 light chain, an antibody fragment used to stabilize the active form of Arr2^23^, at its C-terminus with a 12 amino acid linker (GSAGSAGSAGSA). The construct of the fusion complex was co-expressed with His8-tagged Fab30 heavy chain and a GPCR kinase, GRK5, which phosphorylated the receptor for better arrestin binding. The complex protein was purified using a nickel-affinity column in the presence of 2 µM NTS and 10 µM ML314, then concentrated and further purified by gel filtration chromatography for structural studies (Fig. 1d&e, see methods section for details).

The purified BRIL–NTSR1–Arr2–Fab30 complex protein displayed a relatively high melting temperature (Tm) at 58°C determined by thermostability shift assay^24^ (Fig. 1f). It displayed a uniform distribution when inspected by negative staining EM (Fig. 1g). Single particle cryo-EM data was collected with a FEI Titan Krios microscope. A total number of 17,206 micrographs were recorded, and over 5 million complex particles were picked and sorted for further data processing. 2D classification identified multiple conformations of complex particles. After intensive 3D classification, 220,464 particles (~4.5% of total initial particles) were selected for structure reconstruction (Extended Data Fig. 1). The density map showed an average resolution of 4.8 Å determined by the criterion of Fourier shell correlation (FSC) of 0.143, with the core structure composed of Arr2 and Fab variable region (Fv) showing a resolution range up to 4.5 Å (Fig. 2a& Extended Data Fig. 2). However, the conformational heterogeneity between NTSR1 and Arr2 is still present as illustrated in the multi-body analysis (Extended Data Fig. 3), even though a very low fraction of the full dataset was used in the final reconstruction.

**Figure 2.**
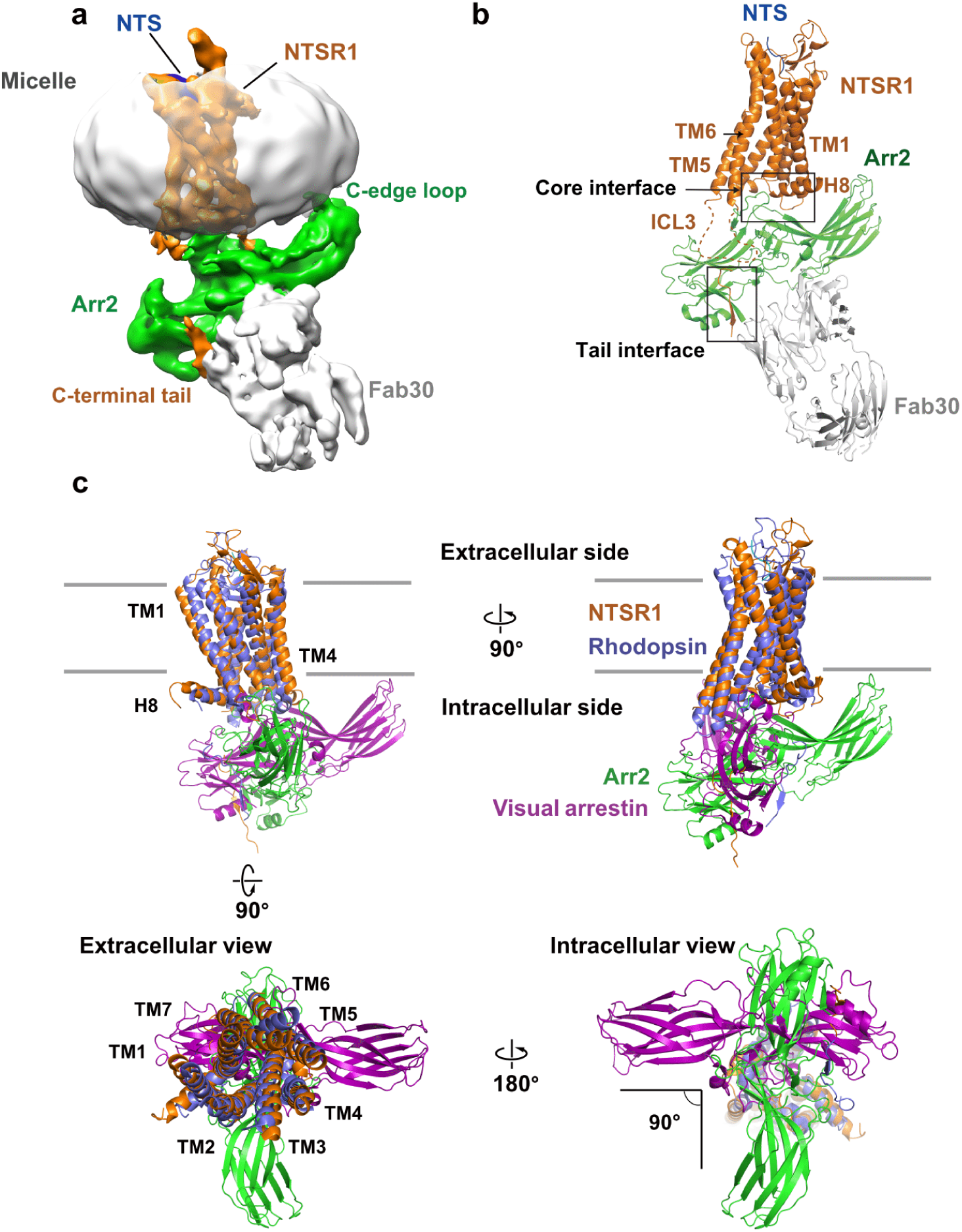
Structural model of Arr2–NTSR1 complex a. Electron density map of NTSR1–Arr2–Fab30 complex structure. While a reasonable contour level for the whole complex structure is about 0.016, the contour level of this map is set to 0.013 to show the whole micelle. NTSR1 is labeled in brown, Arr2 in green, Fab30 in gray, and the micelle surrounding the transmembrane domain of NTSR1 in silver. b. The overall structural model of NTSR1–Arr2–Fab30 complex. Key structural elements are labeled with highlighted boxes for the core interface (including two interface patches) and the tail interface between the receptor and Arr2. The same color code as in panel (a) is used for the components of the complex. c. Superposition of NTSR1 with rhodopsin results in differently oriented core structures of Arr2 and visual arrestin that intersect at a right angle. Four views of the overlaid Arr2–NTSR1 and visual arrestin-rhodopsin complexes with Arr2 colored in green, NTSR1 in brown, visual arrestin in magenta, and rhodopsin in purple. The PDB code of the structure model of rhodopsin-visual arrestin complex is 5W0P.

Despite moderate resolution, the EM map allowed clear determination of the position and orientation of NTSR1, Arr2, and Fab30 in the complex assembly (Fig. 2). The complex model was built by docking the crystal structure of the NTSR1-NTS complex (PDB: 4GRV)^14^, and the structure of Arr2 bound with phosphorylated vasopressin C-terminal peptide (V2Rpp) and Fab30 (PDB: 4JQI)^23^ into the density map, followed by manual model adjustment, real space refinement, and iterative cycles of Rosetta refinement and model rebuilding against the EM map^25^. The final complex model contains NTSR1 residues from R49 to T416, of which ICL3 (A270 to P292) and the linker region between helix 8 and the C-terminal tail (P384 to S404) are missing. The model of Arr2 includes residues T6 through M352 and the model of Fab30 contains a total of 409 residues that comprise both the light chain and the heavy chain of the Fab. The N-terminally fused BRIL and small molecule ML314 are not visible in the density map, and thus not included in the complex model. The statistics and geometry of the structural model are summarized in Extended Data Table 1.

### The overall structure of the Arr2–NTSR1 complex

The Arr2–NTSR1 complex is formed by asymmetric assembly of the two components with the horizontal axis of Arr2 intersecting the inner surface of the cell membrane at about 20°, resulting in the interaction of the hydrophobic C-edge loops (L191 and M192, and L334 through L338) of the arrestin with the cell membrane (Fig 2a&b). This is consistent with the crystal structure of visual arrestin-rhodopsin complex, which for the first time displayed the asymmetric arrestin-GPCR assembly and confirmed the function of the hydrophobic C-edge loops of arrestin as a membrane anchor that stabilizes the binding of arrestin to membrane-embedded receptor, which was later confirmed experimentally^7,26,27^. The interaction of the hydrophobic C-edge loops of the Arr2 with cell membrane layer suggests that the lipid binding capability is a general property of the arrestin family members.

Despite the similarity in the asymmetric assembly of the Arr2–NTSR1 complex with that of the Arr1-rhodopsin complex, the relative orientation of arrestin to the receptor is dramatically different between these two arrestin-receptor complexes. Superposition of the receptor TMDs of these two complexes reveals that the orientation of Arr2 is rotated by 90° around the transmembrane axis from that of visual arrestin (Fig. 2c). In this new orientation, the positions of ICL3 and both TM5 and TM6 of the receptor are pointed toward the N-terminal domain of Arr2, making it possible for ICL3 to resemble the C-terminal tail to interact with the N-domain of Arr2 (Fig. 2b).

To assess the conformational stability of the complex assembly, we performed six independent, two microsecond-long all-atom molecular dynamic simulations of the Arr2–NTSR1 complex without the stabilizing Fab30. Throughout the course of simulation, the C-edge loop L334 through L338 remained in association with the membrane lipids and the NTSR1 C-terminal tail remained in interaction with the N-terminal domain of Arr2 (Extended Data Fig. 4&5). However, the arrestin core structure can rotate around the cryo-EM structure to a limited extent, and the extended finger loop can disengage from the NTSR1 TMD core (Extended Data Fig. 5). These simulation data suggest that the finger loop as well as the core interface between Arr2 and the receptor is highly dynamic and the assembly of Arr2–NTSR1 complex captured by the cryo-EM structure is probably representing one of many conformational states, which is consistent with the fact that the conformational heterogeneity of Arr2-NTSR1 complex assembly still exists in the particles of the 3D class used in the final cryo-EM reconstruction (Extended Data Fig. 3).

### The Arr2-NTSR1 core interface

The Arr2–NTSR1 complex is assembled through intermolecular interactions that consist of two separated major interfaces: a core interface consisting of two separated patches between the central crest loops of arrestin and the intracellular side of the receptor TMD, and a tail interface between the N-terminal domain of Arr2 and the receptor C-terminal tail (Fig. 2b). One patch of the core interface between Arr2 and NTSR1 is the finger loop (from E66 through L71) of the arrestin, which is inserted into the intracellular cavity of the receptor TMD and surrounded by ICL1, ICL2, TM5 and 6, of the receptor (Fig. 3a). In this configuration, the top surface of the finger loop forms a direct interface with the turn between TM7 and helix 8 of the receptor (Fig. 3a).

**Figure 3.**
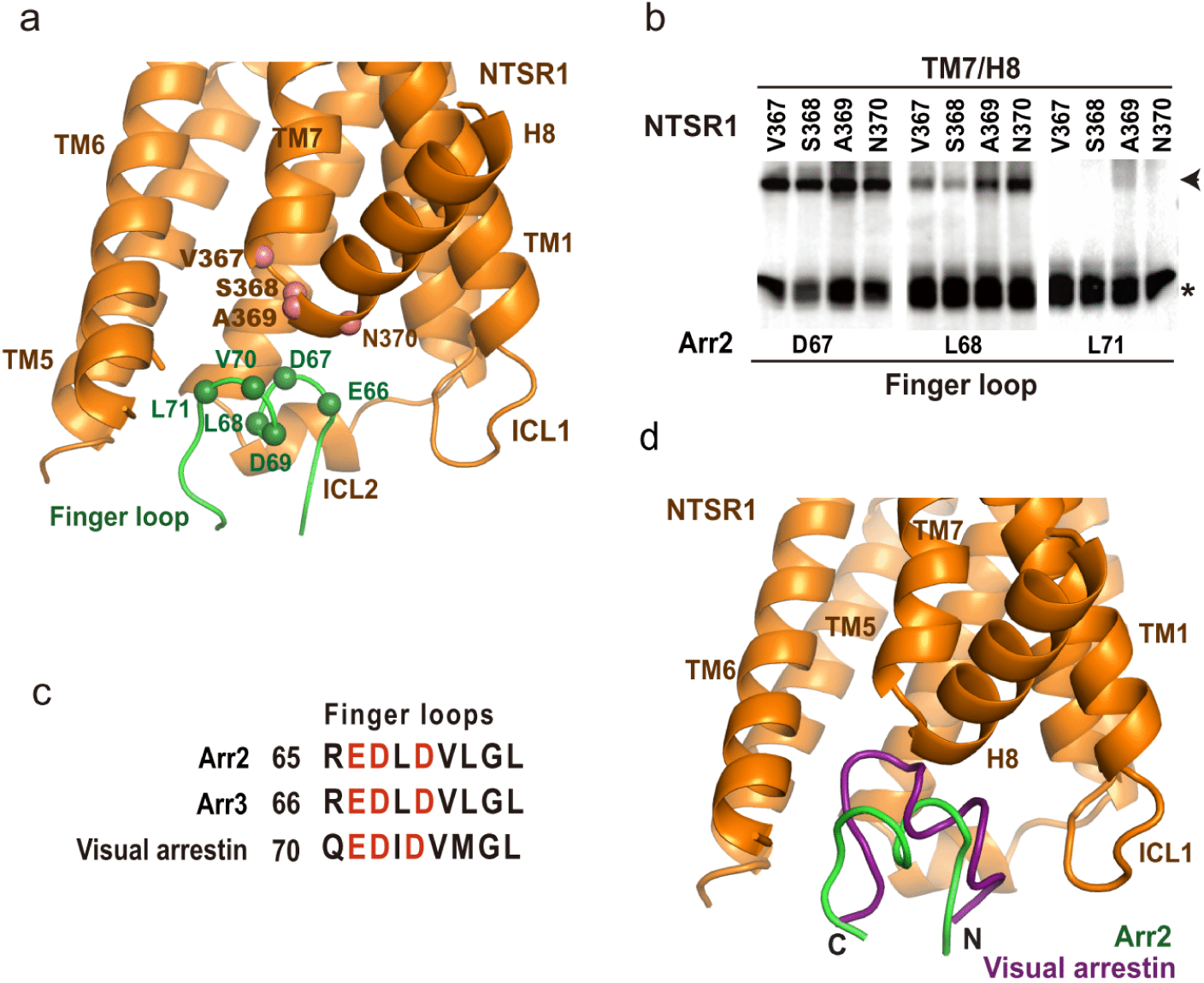
The core interface patch between Arr2 finger loop and NTSR1 TMD a. The interface patch between the finger loop of Arr2 and the intracellular cavity of the receptor TMD. The positions of the major interface residues on arrestin finger loop and the turn between TM7 and H8 of the receptor are indicated by spheres. NTSR1 is colored in brown and Arr2 in green. b. Disulfide crosslinking signals between arrestin finger loop residues D67 and L68, and receptor residues V367, S368, A369 and N370 at the turn between TM7 and helix 8 were strong. As a comparison, no or very weak crosslinking signals were observed between Arr2 residue L71, which is at the C-terminal end of the finger loop. c. Conserved negatively charged residues in the finger loop of Arr2 and Arr3 and visual arrestins d. Superposition of NTSR1 with rhodopsin results in differently oriented core structures (omitted) but well-aligned finger loops of Arr2 and visual arrestin. Rhodopsin is omitted for clarity. Visual arrestin is colored in magenta and Arr2 in green.

To validate the interface of the complex structure, we performed cell-based disulfide crosslinking experiments. In this set of experiments, site-specific cysteine residues were introduced at the interface of Arr2 and NTSR1. Both proteins with cysteine mutations were co-expressed in cells as separate proteins. The crosslinking products were formed by treating cells with oxidizing reagent and were monitored by SDS-PAGE followed by western blotting to detect the Flag tag that was fused to the C-terminus of Arr2. The same method has been used to validate the visual arrestin-rhodopsin complex interface^7,9^. Crosslinking reactions of a total of 177 residue-pairs were performed, among which we observed strong crosslinking signals between Arr2 residues D67 and L68 at the central region of the finger loop, with NTSR1 residues V367 through N370 located at the turn between TM7 and helix 8 (Fig. 3b, and the positions of those interface residues are presented as spheres in Fig. 3a). As a comparison, residue L71 at the C-terminal end of the finger loop showed no or very weak crosslinking signal with those NTSR1 residues (Fig. 3a&b). Several finger loop residues are also found to crosslink with residues at ICL1 (Q98) and TM6 (R294 and A297), which are components of the intracellular cavity of the receptor TMD (Extended Data Fig. 6). These crosslinking data validated the interface patch between Arr2 finger loop and the receptor and showed that the arrestin finger loop serves as a key component of the core interface of this Arr2-GPCR complex.

The amino acid sequences of Arr2 finger loops are enriched with acidic residues, similar to that of visual arrestin (Fig. 3c), which is known to bind to the positively charged TMD cavity^7^. When NTSR1 and rhodopsin are overlaid, the finger loop of Arr2 occupies the same space as the corresponding region of visual arrestin, but it does not insert as deep into the TMD cavity as the visual arrestin (Fig. 3d), indicating that the association of the Arr2 finger loop with the intracellular cavity of NTSR1 TMD could be more dynamic than that between visual arrestin and rhodopsin.

The other patch of the core interface between Arr2 and NTSR1 is the interaction of the crest regions of Arr2 with ICL1 and ICL2 of the receptor, which is different from that in the visual arrestin-rhodopsin complex. In contrast to the similar orientations of the finger loops of the two arrestins, superposition of the two receptors left the core structures of the arrestins in two orientations that differ by an angle of about 90° as shown in Figure 2c. The different orientations of Arr2 and visual arrestin core structures in their GPCR complexes result in different binding modes of the crest regions of arrestins with the intracellular loops of the receptors, particularly ICL1 and ICL2, in these two arrestin-GPCR complexes (Fig. 2c&4).

**Figure 4.**
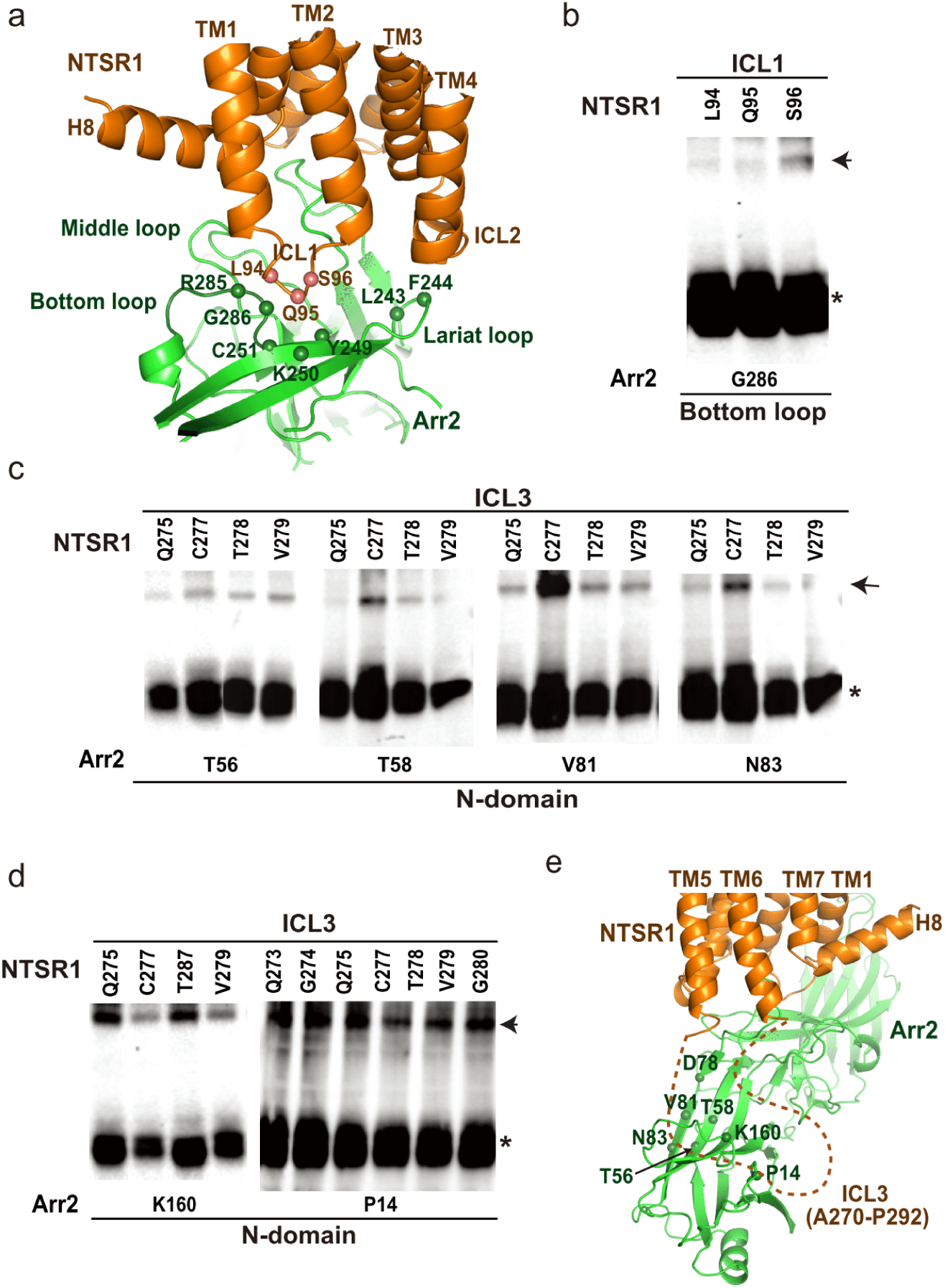
The core interface patch between Arr2 and NTSR1 a. The interface patch between Arr2 and NTSR1, in which ICL1 of NTSR1 interacts with both lariat loop and bottom loop of Arr2. The positions of the interface patch residues are indicated by spheres and validated by disulfide crosslinking shown in panel (b). b. Disulfide crosslinking between bottom loop residue G286 of the receptor with ICL1 residues L94, Q95 and S96 of the receptor. c. Disulfide crosslinking between ICL3 residues of the receptor and residues from the top surface of the Arr2 N-domain. d. Disulfide crosslinking of Arr2 N-domain residues P14 and K160 with ICL3 residues Q273, G274, Q275, C277, T287, V279, G280 and E282. e. The core interface between Arr2 and NTSR1 with a potential interface patch between NTSR1 ICL3 (indicated by dashed line) and the N-domain surface of Arr2 suggested by disulfide crosslinking data shown in panels (c) and (d). The locations of the potential interface residues on N-domain of Arr2 are indicated by spheres. Same color code is used as in panel (a).

Structural comparison of NTSR1 and rhodopsin reveals that they share a conserved TMD conformation with their ICL1 adopting an extended loop and ICL2 forming a short helix in their respective arrestin-bound complexes. The ICL2 helix of rhodopsin is inserted into the cleft formed by the middle loop, bottom loop, and lariat loop of visual arrestin (termed MBL cleft)^7^. However, the same MBL cleft in Arr2 is occupied by the ICL1 of NTSR1 (Fig. 4a) because of the different orientation of the arrestin. Correspondingly, ICL2 of NTSR1 is rotated away from the MBL cleft to reposition to the top of the lariat loop (Fig. 4a). Disulfide crosslinking experiments showed crosslinking signals between receptor residues L94 through S96 on ICL1 of NTSR1 and residue G286 at the bottom loop of Arr2, which is consistent with the interface of ICL1 of NTSR1 with this crest region of Arr2 (Fig 4b).

ICL3 of the receptor, despite being invisible in the density map, likely forms an interface with the N-domain of β-arrestin based on the Arr2–NTSR1 complex conformation (Fig. 2b). Disulfide crosslinking data showed that ICL3 residues Q275, C277, T278 and V279 of NTSR1 crosslinked with residues T56, T58, V81 and N83 at the top surface of the Arr2 N-domain (Fig. 4c & Extended Data Fig. 6). These ICL3 residues also show strong crosslinking signals with Arr2 residue K160 (Fig. 4d). Additional disulfide crosslinking signals were observed between most residues from Q273 through G280 of the NTSR1 ICL3 and residue P14 from Arr2 (Fig. 4d). These crosslinking data indicate that ICL3 of the receptor is positioned close to the N-domain and may interact with residues at the N-domain surface of Arr2 (Fig. 4e).

### The Arr2–NTSR1 tail interface

While the Arr2–NTSR1 core interface is highly dynamic, the tail interface between NTSR1 C-terminal tail and the Arr2 N-domain seems similar to that in the visual arrestin-rhodopsin complex^9^ (Fig. 5). We observed electron density at a contour level of 0.016 (at which the density of the overall complex structure can be properly shown) of about ten C-terminal tail residues of NTSR1 that bind at the first β-strand of Arr2 (Fig. 5a). However, detailed information about this region of the C-terminal tail, including its specific residues, is difficult to identify due to the limited resolution of the density map. Analyses of NTSR1–Arr2–Fab30 fusion protein by mass spectrometry (Fig. 5b & Extended Data Fig. 7) revealed that seven serine or threonine residues from S401 through T416 on NTSR1 C-terminal tail were phosphorylated, suggesting possible involvement of this region in binding to the positively charged surface of the arrestin N-domain. Disulfide crosslinking experiments showed that Arr2 residue P14 at the C-terminal turn of the first β-strand strongly crosslinked with receptor residues H406 through S410 (Fig. 5d & Extended Data Fig. 6), which indicated that receptor residues C-terminal to H406 are likely the region that binds to arrestin N-domain (Fig. 5c)

**Figure 5.**
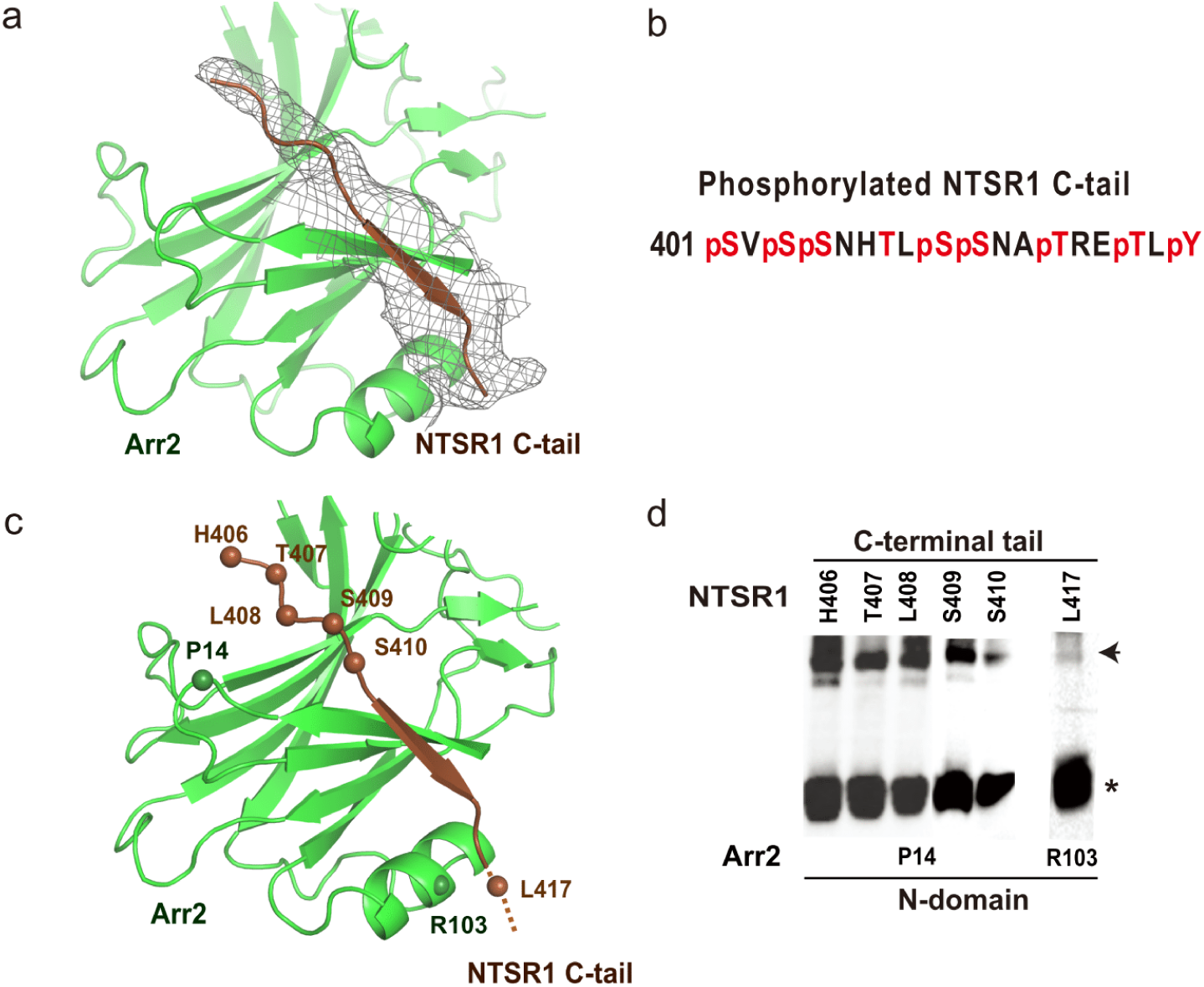
The interface between a C-terminal tail of NTSR1 and the positively charged N-domain of Arr2 a. An overall EM map of the NTSR1 C-terminal tail that binds at the first β-strand of Arr2. The density map at contour level of 0.016 covers about ten C-terminal tail residues of the receptor. The receptor C-terminal tail is colored in brown, Arr2 in green, and EM map in gray. b. Mass spectrometry analysis of NTSR1–Arr2–Fab30 fusion protein revealed that C-terminal residues S401, S403, S404, S409, S410, T413, T416 and Y418 are phosphorylated in the protein sample. c. The tail interface between the NTSR1 C-terminal tail and the first β-strand of Arr2. Disulfide crosslinking (shown in panel d) suggests that the region of NTSR1 C-terminal tail that binds to the Arr2 N-domain is likely in between residues H406 and L417. The residues that show crosslinking signals are labeled based on crosslinking data in panel (d). d. Disulfide crosslinking data of P14, at the top surface of the Arr2 N-domain, with the receptor C-terminal residues H406 through S410; and residue R103 of β-arrestin crosslinked with receptor C-tail residue L417.

### A model for Arr3-GPCR interaction

The human genome contains two subtypes of β-arrestin: Arr2 and Arr3, which share over 53% sequence identity but have both some overlapping and some divergent functions. As Arr3 showed NTSR1 binding affinity similar to that of Arr2 (Fig. 1b), we wanted to test whether the binding mode of Arr3 to NTSR1 is conserved with that of Arr2. Our disulfide crosslinking experiments displayed that Arr3 finger loop residues E67 through L72 (corresponding to E66 through L71 of Arr2) crosslinked with the corresponding set of residues (A369 and N370) at the turn between TM7 and helix 8 of NTSR1 (Extended Data Fig. 8), indicating a conserved finger loop interface with TM7 and H8 of the receptor between these two β-arrestins.

In addition, disulfide crosslinking data shows that Arr3 residue P15 (corresponding to P14 of Arr2) crosslinked with residues N405 through T407 of the receptor C-terminal tail, suggesting a similar tail interface between Arr3 and NTSR1 (Extended Data Fig. 8). Similar crosslinking pattern between NTSR1 ICL3 residues and Arr3 N-domain residues, including P15 at the C-terminus of the first β-stand and K161 (corresponding to K160 of Arr2) on the loop between the top β-strands were also observed (Extended Data Fig. 8). Together, these crosslinking data suggest that the overall assembly of Arr3 with NTSR1 is similar to that of Arr2 to NTSR1 in this core engaged configuration.

### A model for β-arrestin recruitment by 5-HTR1A and 5-HTR1B

Many GPCRs, including serotonin receptors 5-HTR1A and 5-HTR1B as well as several dopamine receptors, either lack or contain a very short C-terminal tail and they are proposed to use their ICL3 as an alternative interface to recruit β-arrestins^28^. 5-HTR1A and 1B have long ICL3s with multiple phosphorylation sites for potential interaction with positively charged N-domains of β-arrestins. Our biochemical binding assays demonstrated that both receptors bind to Arr2 and 3 with similar activity to that of NTSR1 with Arr2 (Extended Data Fig. 9). However, when the visual arrestin–rhodopsin complex structure was used as a template, it was difficult to model the binding mode of the phosphorylated ICL3 of 5-HTR1A or 1B to the positively charged cleft on the N-domain of β-arrestins. That is because TM5 and 6 as well as ICL3 of rhodopsin are positioned at the opposite site to the positively charged surface of the arrestin N-domain in the visual arrestin–rhodopsin complex structure.

The Arr2–NTSR1 complex structure, however, displays a distinct complex assembly with Arr2 rotated by 90° from the position of visual arrestin in the arrestin-rhodopsin-complex (Fig. 2c). TM5 and TM6 of NTSR1 are, therefore, positioned above the front surface of the N-domain of Arr2, allowing ICL3 of the receptor (not visible in the structure) to reach the positively charged N-domain of Arr2 (Fig. 4e). Our disulfide crosslinking data provide evidence that NTSR1 ICL3 can interact with the positively charged cleft of the N-domain of Arr2 and 3 (Fig. 4c-e, Extended Data Fig. 6&8). Therefore, the structure of the Arr2–NTSR1 complex may serve as a suitable template to model the interface of β-arrestins with 5-HTR1A and 5-HTR1B.

Using the Arr2-NTSR1 structure as a template, and the crystal structure of ergotamine-bound 5-HTR1B as an initial model for 5-HTR1B (PDB: 4IAR)^29^, we generated an Arr2–5-HTR1B complex homology model, in which the core interface between Arr2 and 5-HTR1B is similar to that of Arr2-NTSR1 complex (Fig. 6a&b). We then performed disulfide crosslinking experiments to test the complex assembly of Arr2–5-HTR1B shown in the homology model (Fig. 6c & Extended Data Fig. 10).

**Figure 6.**
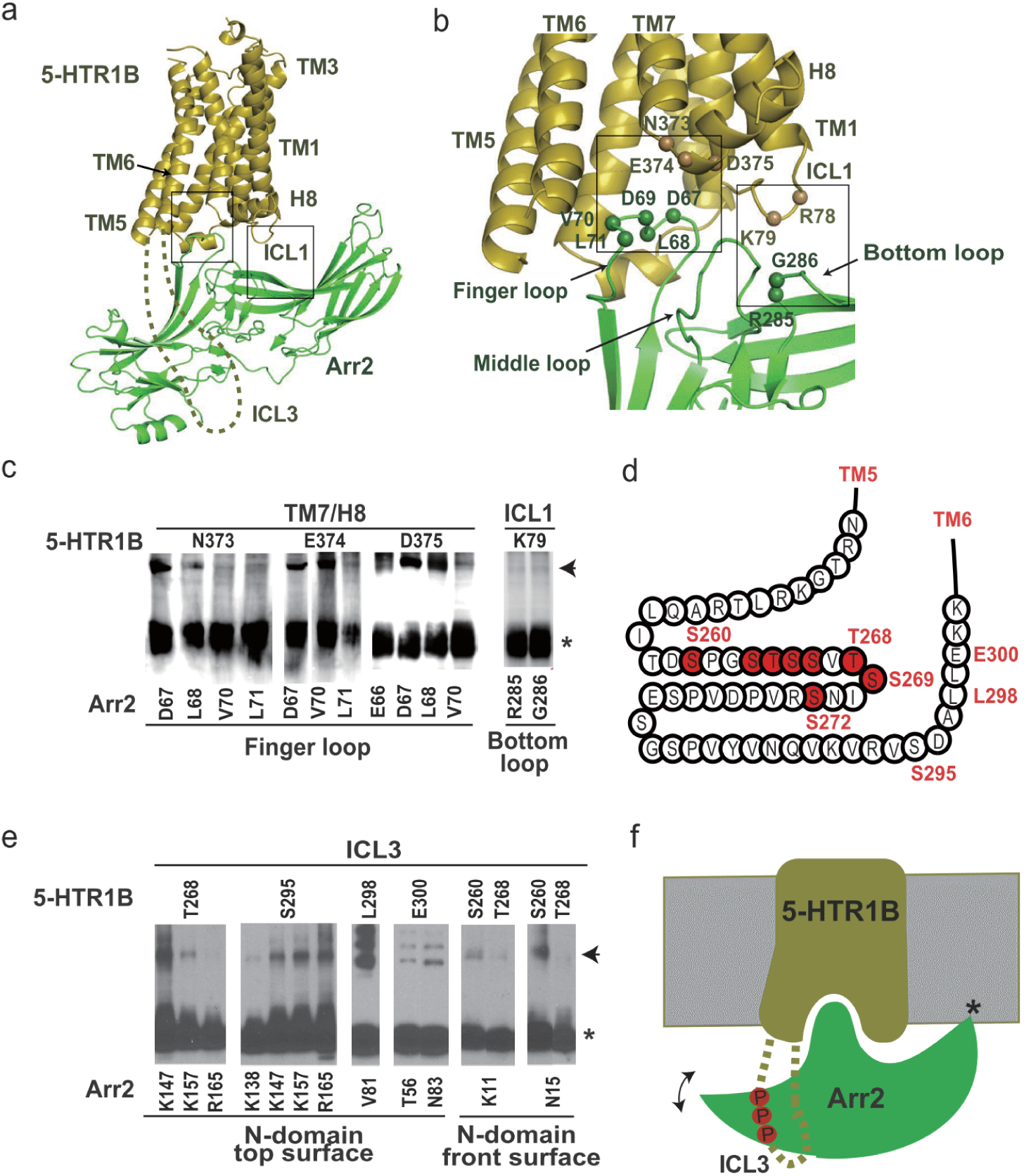
The binding mode of Arr2 with 5-HTR1B a. Arr2–5-HTR1B homology model based on Arr2–NTSR1 complex as a template. Highlighted in boxes are core interface patches between Arr2 and 5-HTR1B. b. Core interface patches between Arr2 and 5HTR1B with interface residues (shown as spheres) validated by disulfide crosslinking shown in panel (c). c. Disulfide crosslinking between residue pairs at Arr3–5-HTR1B core interface patches. d. A cartoon presentation of the 5-HTR1B ICL3 extended from TM5 and TM6 with phosphorylation code residues highlighted in red. e. Disulfide crosslinking between residue pairs at the interface between Arr2 N-domain and 5-HTR1B ICL3. f. A cartoon model to illustrate the interaction between ICL3 of 5-HTR1B with the positively charged N-domain of Arr2. The model indicates the interaction of arrestin with the receptor TMD core and ICL3, as well as the lipid anchoring of the arrestin C-edge loops (indicated by a star). The double-arrow indicates the conformational swing of arrestin around the receptor core.

We observed crosslinking signals between finger loop residues D67 L68, V70, and L71 of Arr2 and residues N373, E374 and D375 at the turn between TM7 and helix 8 of 5-HTR1B (Fig. 6c). Residues D67, L68, V70, and L71 of the finger loop crosslinked with receptor residues R308 and K311 at the inner surface of TM6 (Extended Data Fig. 10). These crosslinking data support a conserved binding mode of the Arr2 finger loop with the intracellular cavity of the 5-HTR1B TM domain (Fig. 6a&b). Disulfide crosslinking data also showed crosslinking signals between Arr2 residues R285 and G286 in the arrestin bottom loop with ICL1 residue K79 of 5-HTR1B (Fig. 6c & Extended Data Fig. 10). This specific crosslinking is only consistent with the arrestin orientation in the Arr2-NTSR1 model, but it is not compatible with the visual arrestin-rhodopsin complex. Together these crosslinking data support a core interface between the Arr2 crest region and the intracellular side of the 5-HTR1B TM domain, which is conserved with that in the Arr2–NTSR1 complex (Fig. 6a-c).

5-HTR1B contains multiple serine and threonine residues in the central area of its long ICL3 that may be phosphorylated for binding to the surface of the positively charged arrestin N-domain (Fig. 6d). We designed cysteine pair mutations in the potential interface between ICL3 of the receptor and the N-domain of Arr2 and found that ICL3 residues T268, S295, L298 and E300 were crosslinked with residues at the surface of the top β-sheet, including T56, V81, N83, K147 and K157 of Arr2 (Fig. 6e & Extended Data Fig. 10). Residues S260 and T268 on ICL3 showed crosslinking signals with residues K11 on the first β-strand and N15 on the turn following the first β-strand of Arr2 (Fig. 6e & Extended Data Fig. 10). Residue S260 is near the C-terminal end of TM5, and its crosslinking with K11 at the positively charged side of the N-domain of Arr2 is only possible in the Arr2-NTSR1 orientation but not in the visual arrestin-rhodopsin orientation because in the visual arrestin–rhodopsin complex structure, both TM5 and 6 of the receptor are positioned at the back site of the positively charged surface of the arrestin N-domain.

We also observed a nearly identical Arr2 crosslinking pattern between 5HTR1A and 5-HTR1B (see supplemental text). These crosslinking data, together with the homology model of Arr2–5-HTR1B complex, have provided an overall picture of how 5-HTR1A and 5-HTR1B recruit Arr2. While the intracellular core interface between the TM domain of 5-HTR1A and 5-HTR1B with Arr2 is similar to that in the Arr2-NTSR1 complex, the ICL3 of 5-HTR1A and 5-HTR1B can extend down from the intracellular sides of TM5 and TM6, with their phosphorylated serine or threonine residues forming charge interactions with the positively charged surface on the Arr2 N-domain, forming an interface that resembles the tail interface in the Arr2–NTSR1 complex (Fig. 6a).

## Conclusions

In this paper, we report a structure of Arr2 in complex with NTSR1, a class A GPCR, by cryo-EM single particle analysis. While the binding mode of the tail interface between the C-terminal tail of the receptor and the positively charged arrestin N-domain is conserved with that in the visual arrestin-rhodopsin complex, this structure displays a different Arr2 orientation from that seen in the visual arrestin-rhodopsin complex. Homology modelling and disulfide crosslinking interface mapping demonstrate that this core interface is conserved in Arr2 complexes with 5-HTR1A and 1B subfamily. In these complex models, the orientation of Arr2 allows ICL3 of the receptor to reach the surface of the Arr2 N-domain, providing the possibility for a GPCR that lacks a C-terminal tail to use its ICL3 to interact with the arrestin N-domain. As Arr2 and Arr3 are ubiquitously expressed protein partners for signal transduction of many nonvisual GPCRs, our structural and biochemical studies of Arr2 interaction with NTSR1 and 5-HTR1 subfamily members have provided an alternative model for understanding the interaction of Arr2 and Arr3 with nonvisual GPCRs.

### Supplementary text on A model for β-arrestin recruitment by 5-HTR1A

We assume that 5-HTR1A shares a conserved receptor conformation with 5-HTR1B and forms a complex structure with Arr2 similar to that of 5-HTR1B, despite that no 5-HTR1A 3D structure has been reported. To test this assumption, we designed corresponding sets of cysteine mutations and performed disulfide crosslinking experiments with the same residues as used to validate Arr2–5-HTR1B complex assembly. We observed crosslinking signals between finger loop residues D67, L68, V70 and L71 of Arr2 and residues N404, K405 and D406 (corresponding to N373, E374, and D375 of 5-HTR1B) at the turn between TM7 and helix 8 of 5-HTR1A (Extended Data Fig. 11). Residues D67 and L68 of the finger loop were also observed to crosslink with receptor residues R339 and K342 (corresponding to R308 and K311 of 5-HTR1B) at the inner surface of TM6 (Extended Data Fig. 11). We further observed disulfide crosslinking between Arr2 bottom loop residues R285 and G286 and ICL1 residue S66 of 5-HTR1A (corresponding to K79 of 5-HTR1B) (Extended Data Fig. 11). These crosslinking data support a similar core interface between Arr2 with 5-HTR1A and 5-HTR1B, which is conserved with that in the Arr2–NTSR1 complex (Fig. 6a&c).

The ICL3 of 5-HTR1A contains more than 100 amino acid residues (233 through 336) with 12 serine or threonine residues that constitute six partial phosphorylation codes together with glutamic acid or aspartic acid residues (Extended Data Fig. 12). We designed cysteine pair mutations to map the potential interface between ICL3 of this receptor and the N-domain of Arr2. Crosslinking data displayed that ICL3 residues V230, T240, P250, R261, K270 and D285 crosslinked with both K160 on a loop connecting the top β-strands of the Arr2 N-domain (Extended Data Fig. 11). Residue P14 on the C-terminal end of the first β-strand of Arr2 strongly crosslinked with ICL3 residues N300, P308, P315, P318, P321 and R323 of 5-HTR1A (Extended Data Fig. 11). While these crosslinking data support a binding interface between ICL3 of 5-HTR1A with Arr2 N-domain, its binding mode is likely different from that of 5-HTR1B ICL3 with Arr2 due to the largely different sequences between the two ICL3s of these two serotonin receptors.

## Methods

### Constructs and expression of BRIL–NTSR1–Arr2–Fab30 complex

In this study, we used human neurotensin receptor 1 (UniProtKB ID: P30989, NTSR1 residues 49–418) and human Arr2 with 3A mutations (UniProtKB ID: P49407, residues 1–393, with mutations I386A, V387A, F388A). To obtain a stabilized complex, we used a fusion construct of BRIL–NTSR1–Arr2–Fab30L generated by fusing Arr2 to the C-terminus of NTSR1 with a 3 amino acid linker (GSA), and fusing the light chain of antibody fragment Fab30 to C-terminus of Arr2 with a 12 amino acid linker (GSAGSAGSAGSA). The cytochrome b562 RIL domain (BRIL) was fused at the N-terminus of the receptor. Additional mutations were introduced for the purpose to obtain more stable complex: V81C in Arr2 to form a disulfide bond to the residue C277 in the ICL3 of NTSR1, A279C in Arr2 and G59C in the heavy chain of antibody fragment Fab30 to form a disulfide bond. The BRIL-NTSR1–Arr2–Fab30L fusion construct was cloned into pFastBac (Invitrogen) baculovirus expression vector with a haemagglutinin (HA) signal sequence. The heavy chain of antibody fragment Fab30 was cloned into pFastbac vector with 8xHis tag at the C-terminus. Human GRK5 (UniProtKB ID: P34947) with three mutations, Q41L, K454A and R455A, was cloned into a pFastBac vector. All genes were codon optimized and synthesized by Genewiz for insect cell expression, and all mutants were generated using the Mut Express MultiS Fast Mutagenesis Kit V2 (Vazyme Biotech Co.,Ltd) and verified by DNA sequencing. All constructs were expressed in *Spodoptera frugiperda* (Sf9) cells using baculovirus. Cell cultures were grown in ESF 921 serum-free medium (Expression Systems) to a density of 2-3 million cells per ml and then infected with three separate baculoviruses at a ratio of 2:2:1 for BRIL-NTSR1–Arr2–Fab30L, Fab30H-His8 and GRK5 at a multiplicity of infection (m.o.i.) of about 5. The cells were collected by centrifugation 48 h after infection at 27 °C and cell pellets were stored at − 80 °C until use.

### BRIL–NTSR1–Arr2–Fab30 complex purification

Insect cell membranes were disrupted by thawing frozen cell pellets in a hypotonic buffer containing 10 mM HEPES at pH 7.5, 10 mM MgCl_2_, 20 mM KCl and EDTA-free protease inhibitor cocktail (TargetMol), followed by Dounce homogenization. The lysate was centrifuged at 100,000×g at 4 °C for 1 h. Extensive washing of the raw membranes was performed by repeated Dounce homogenization in a high osmotic buffer containing 1.0 M NaCl, 10 mM HEPES at pH 7.5, 10 mM MgCl_2_, 20 mM KCl, and EDTA-free protease inhibitor cocktail (Roche?) (2 times). By extensive washing, soluble and membrane associated proteins were separated from integral transmembrane proteins.

Washed membranes were solubilized in 20 mM HEPES (pH 7.5), 200 mM NaCl, 25 mM imidazole, 10% (v/v) glycerol, 5 µM peptide NTS (8-13) (synthesized by Synpeptide Co, Ltd) 0.5% (w/v) n-dodecyl-β-D-maltopyranoside (DDM, Anatrace) and 0.1% (w/v) cholesteryl hemisuccinate (CHS, Anatrace) plus EDTA-free protease inhibitor cocktail at 4 °C for 3-4 h. The supernatant was isolated by centrifugation at 100,000×g for 30 min, followed by incubation with Ni-NTA beads (GE Healthcare) for 2 h at 4 °C. After binding, the beads were washed with 10 column volumes of Washing A Buffer (20 mM HEPES, pH 7.5, 200 mM NaCl, 25 mM imidazole, 2 µM NTS(8-13), 10% (v/v) glycerol, 0.05% (w/v) DDM, 0.01% (w/v) CHS and 10 mM ATP), followed by 5 column volumes of Washing B Buffer (20 mM HEPES, pH 7.5, 200 mM NaCl, 50 mM imidazole, 2 µM NTS(8-13), 10% (v/v) glycerol, 0.1% (w/v) digitonin (Anatrace)). The protein was eluted with 3-4 column volumes of Elution Buffer (20 mM HEPES, pH 7.5, 200 mM NaCl, 300 mM imidazole, 2 µM NTS (8-13), 0.1% (w/v) digitonin). The BRIL-NTSR1-Arr2-Fab30 complex sample was concentrated with a 100 kDa molecular weight cut-off Centrifugal Filter (Millipore Corporation) and then size-separated onto Superdex S200 10/300 GL column in 10 mM Tris, pH 7.4, 100 mM NaCl, 0.1% digitonin, 2 µM NTS (8-13) and 10 µM ML314. The fractions for the monomeric complex were collected and concentrated for EM experiments. Typically, the protein yield is about 1-2 mg per 2 L cell culture.

### Cryo-EM sample preparation and data acquisition

The cryo-EM samples were prepared by applying an aliquot of 3 μL protein sample of BRIL–NTSR1–Arr2–Fab30 complex (7 mg/mL) to a glow-discharged Quantifoil holey carbon grid, blotted with filter paper for 2.0 s and plunge-frozen in liquid ethane using an FEI Vitrobot Mark IV. Cryo-EM micrographs were collected on a 300 kV Titan Krios microscope (FEI) equipped with a Gatan energy filter (operated with a slit width of 20 eV) (GIF) and K2 Summit direct detection camera. The microscope was operated at a calibrated magnification of 48,497 Х, yielding a pixel size of 1.031 Å on micrographs. In total, 17,206 micrographs were collected at an electron dose rate of ~8.5 e^−^/Å^2^•s with a defocus range of −2.0 μm to −3.5 μm. An accumulated dose of 68 e^−^/Å^2^ on sample was fractionated into a movie stack of 32 image frames.

### Image processing

For each movie stack, the frames were aligned for beam-induced motion correction using the program MotionCor2^30^. Gctf^31^ was used to determine the contrast transfer function parameters. About 3,000 particles were initially picked manually from selected micrographs, and then subjected to a round of reference-free 2D classification. Averages representing projections of BRIL–NTSR1–Arr2–Fab30 in different orientations were selected and served as templates for automatic particle picking from the full dataset of 14,058 micrographs. Approximately 5.0 million particles were extracted and subjected to rounds of iterative 2D classification. An initial model was generated via Relion 3.0^32^ and used as a starting map for 3D classifications. For the first round of 3D classification, one of the 3D classes showed a complete overall configuration of the receptor-arrestin-Fab30 complex, including BRIL, receptor, Arr2 and Fab30. Particles from this 3D class were then selected for further 3D classification by many rounds. Finally, 260,322 particles from a 3D average showing good secondary structural features in the transmembrane domain were selected for further 3D refinement, which yield a 4.9 Å reconstruction. 3D-FSC and orientation distribution analysis revealed that this reconstruction had the anisotropic problem, which was resulted from that only a relatively small fraction of top view particles was included in the above reconstruction. To overcome this problem, we added more data from 2D classifications, which contained more top view particles, and carried out an additional round of 3D-classification and auto-refinement. The new reconstruction of the Arr2-NTSR1 complex shows a similar overall resolution (4.8 Å) as the previous one but with improved map quality, and the issue of directional resolution anisotropy was also alleviated Fourier shell correlation at 0.143 was used to estimate the overall resolution of the final reconstruction using the Post-processing function of Relion 3.0 and the local resolution was calculated by ResMap^33^.

### Model building and structure refinement

The complex model was built by docking the crystal structure of the NTSR1-NTS (8-13) complex (PDB: 4GRV)^14^, and the structure of Arr2 in complex with phosphorylated vasopressin C-terminal peptide (V2Rpp) and Fab30 (PDB: 4JQI)^23^ into the density map using UCSF Chimera^34^, followed by manual model adjustment in COOT^35^, and real space refinement using PHENIX programs^36^. While the density map nicely defined most backbone traces of the complex as well as some large side chains of the complex, most side chain rotamers, particularly those at the interface of the complex, are defined by ROSETTA model refinement and rebuilding^37^ against the EM map. The structural model has been carefully inspected, and all key Arr2-NTSR1 interfaces shown in the model have been validated by disulfide crosslinking assays. The model statistics were calculated with MolProbity^38^ and listed in Extended Data Table 1. Structural figures were prepared in Chimera or PyMOL (https://pymol.org/2/).

### In-cell disulfide bond cross-linking

The open reading frames of 3A mutant Arr2 were cloned into pcDNA6 with pre-inserted coding regions for C-terminal 3xFlag; concurrently receptors including full-length NTSR1, 5-HTR1A or 5-HTR1B were amplified and cloned into pcDNA6 with pre-inserted coding regions for C-terminal Hemagglutinin (3xHA) tags, respectively. Single cysteine mutations were designed to probe the receptor/arrestin interface based upon structural and sequence alignments, and systematically introduced into these corresponding DNA vectors. A total of 132 mutants of Arr2 and Arr3, and mutants of NTSR1, 5-HTR1A, and 5-HTR1B, were designed and appropriate combinations of β-arrestin-receptor mutation pairs were tested for disulfide crosslinking. For in-cell disulfide crosslinking, AD 293 cells were split at 50,000 cells per well in a 24-well plate. Cells were grown for one day, then transfected with 100 ng β-arrestin_3A plasmid (pcDNA6-Arr2/2_3A-3Flag) plus 100 ng corresponding receptor constructs (pcDNA6-NTSR1/5-HTR1A/5-HTR1B-3HA) by Lipofectamine 2000 (Invitrogen, DNA/Lipofectamine 2000 ratio of 1:2) in each well. The transfected cells were grown for 48 hours. 10 min prior to oxidation treatment, the receptors were activated by addition of corresponding agonist (5 µM of NTS (8-13) for NTSR1, and 10 µM of 5-HT for 5-HTR1A/5-HTR1B). To induce disulfide bond formation, the cells were incubated for 5 min at room temperature with H_2_O_2_ solution freshly diluted in culture medium to a final concentration of 1 mM. The cells were lysed by 10-minute shaking with 100 µl of CelLytic M (Sigma, C2978) per well. The cell lysates were separated from lysed cells by centrifugation at 16,000×g at 4 °C for 15 min. 12 µl supernatants of each sample were mixed for 5 minutes with 3 µl 5×SDS loading buffer without reducing agent, separated by SDS PAGE, and analyzed by Western blotting using horseradish peroxidase-conjugated goat-anti-mouse (Beyotime Biotechnology, A0216) and mouse anti-Flag (Beyotime Biotechnology, AF519) primary antibodies to detect the expression of free or cross-linked β-arrestin.

### Tango Assay

The cDNA of NTSR1 (1-418, wild type) was cloned into pcDNA6 vector consisting of an expression cassette with tobacco etch virus (TEV) protease cleavage site and the transcriptional activator tTA at the C terminus (pcDNA6-NTSR1-TEV site-tTA). A TEV protease cDNA was fused to the C-terminus of Arr2(1-393, wild type) (pcDNA-Arr2-TEV protease). Interaction between NTSR1 and Arr2 leads to the cleavage of the TEV site, thus releasing tTA to trigger tTA-dependent luciferase reporter gene expression. For Tango assays, HTL cells were cultured in 24-well plate at a density of 5×10^4^ cells/well for 24 h, and then transfected with 10 ng NTSR1-TEV site-tTA, 10 ng Arr2-TEV protease plasmids and 5 ng of phRG-tk Renilla luciferase expression plasmids using X-tremeGENE 9 DNA Transfection Reagent (Roche). After transfection for 24 h, cells were incubated overnight with PBS (vehicle), 5 μM NTS, 5 μM ML314, and 5 μM NTS combined with 5 μM ML314, respectively. Then luciferase activities were evaluated according to manufacturer’s protocols of the Dual Luciferase Kit (Promega). Tango Assays for NTSR1 and the Arr2 3A mutant were carried out the same as above.

### NanoBiT Assay

The cDNA of NTSR1(1-418, wild type) was inserted into the NanoBiT PPI plasmid (Promega) with the large NanoBiT subunit (LgBiT) at its C-terminus (pBiT-NTSR1-LgBiT). The small NanoBiT subunit (SmBiT) was positioned at the N-terminus of Arr2 (1-393, wild type) (pBiT-SmBiT-Arr2). Interaction between NTSR1 and Arr2 brings LgBiT and SmBiT into proximity to form a functional enzyme that can convert NanoLuc substrate to generate a luminescent signal. For NanoBiT Assay, AD293 cells were transfected following the manufacturer’s protocol (FuGENE^®^ HD, Promega). Specifically, AD293 cells were cultured in 6-well plate at a density of 5×10^5^ cells/well for 24 h and then transfected with 1.5 μg pBiT-NTSR1-LgBiT and 1.5 μg pBiT-SmBiT-Arr2 plasmids for 24 h. The transfected cells were harvested and diluted to 4×10^4^ cells/ml using Opti-MEM (Invitrogen) and then plated in 384-well white plates (Greiner) with 20 μl/well. 10 μl/well NanoLuc substrate and 10 μl/well NTS_8-13_ were added before luminescence was detected. NanoBit assays for other combinations including NTSR1 with Arr2 (1-393, 3A), NTSR1 with Arr3 (1-393, wild type) and NTSR1 with Arr3 (1-393, 3A) are the same as above. The NanoBiT assays for 5-HTR1A/5-HTR1B with Arr2 (wild type or 3A) or Arr3 (wild type or 3A) were performed with different concentrations of 5-HT.

### System preparation and molecular dynamics simulation

All-atom atmospheric molecular dynamics simulations of Arr2-NTSR1 complex were performed using the CHARMM36m forcefield ^39^ with the GPU accelerated Particle-Mesh Ewald molecular dynamics (pmemd.cuda) engine within AMBER18 ^40^. The Arr2–NTSR1 complex structure was aligned for membrane insertion using the Orientations of Proteins in Membranes database^41^. Missing residues 384–404 in the C-tail were modelled and subjected to 500 rounds of very slow loop refinement assessed by DOPE scoring using Modeller9.13^42^. The receptor arrestin complex was inserted into a pre-equilibrated POPC lipid bilayer solvated in a box of TIP3P waters with 150 mM NaCl and neutralized by removing appropriate ions or counter ions using the Desmond system builder within Maestro (Schrödinger Release 2018-1: Maestro, Schrödinger). Titratable residues were left in their dominant state at pH 7.0 and all histidine side chains were represented with a hydrogen atom on the epsilon nitrogen. Free protein amino and carboxyl groups were capped with neutral acetyl and methylamine groups. Representative system initial system dimensions were 118 x 118 x 150 Å and comprised of 313 lipids, 39,278 water molecules, 216 sodium ions and 258 chloride ions for a total of approximately 171,000 atoms.

Prior to production simulations, 25,000 steps of energy minimization were carried out followed by equilibration in the canonical NVT and isothermal-isobaric NPT ensembles for 10 and 50 ns respectively with harmonic restraints (10 kcal mol^−1^ Å^−2^) placed on all Cα atoms. Each system was then simulated for an additional 50 ns without harmonic restraints. Production simulations were performed with a 2 fs time-step in the NPT ensemble with semi-isotropic coupling at 310 K and 1 bar maintained by the Langevin thermostat and Monte Carlo barostat with periodic-boundary conditions. Bonds involving hydrogen atoms were constrained by SHAKE and with non-bonded interactions cut at 8 Å. Trajectory snapshots were saved every 10 ps. Simulation analysis was performed using MDTraj 1.7.2^43^ and VMD 1.9.2^44^ and CPPTRAJ^45^. Plots were generated using the R statistical package (https://stat.ethz.ch/pipermail/r-help/2008-May/161481.html).

### Identification of NTSR1 phosphorylation sites using liquid chromatography-mass spectrometry

Protein was digested with trypsin or chymotrypsin and then analyzed on the Easy nano-LC1000 system (Thermo Fisher Scientific) using a self-packed column (75 μm × 150 mm; 3 μm ReproSil-Pur C18 beads, 120 Å, Dr. Maisch GmbH, Ammerbuch, Germany) at a flow rate of 300 nL/min. The peptides were separated across a gradient (2-27% mobile phase B) over a 90 min or 120 min period and analyzed online on either a Q-Exactive or Orbitrap Fusion mass spectrometer (Thermo Fisher Scientific).

The Orbitrap Fusion mass spectrometer was operated in data-dependent mode with each full MS scan (m/z 350 - 1500) followed by MS/MS in 3 s of cycle time. Both HCD and three consecutive scans with CID, ETD and EThcD were acquired on the same precursor. HCD was carried out at 30 normalized collision energy (NCE). CID was carried out at 35 NCE in the ion trap. ETD was performed with calibrated charge dependent reaction time. EThcD was performed with user defined charge dependent reaction time supplemented by 30 NCE HCD activation. Dynamic Exclusion™ was set for 45 s. The full mass and the subsequent MS/MS analyses were scanned in the Orbitrap analyzer with R=60,000 and R=15,000, respectively.

The Q-Exactive mass spectrometer was operated in data-dependent mode with each full MS scan (m/z 350 - 1500) followed by MS/MS for the 15 most intense ions with the parameters: ≥+2 precursor ion charge, 2 Da precursor ion isolation window and 27 NCE of HCD. Dynamic Exclusion™ was set for 30 s. The full mass and the subsequent MS/MS analyses were scanned in the Orbitrap analyzer with R=70,000 and R=17,500, respectively.

The MS/MS spectra were processed using Proteome Discoverer (Version 2.2, Thermo Fischer Scientific, San Jose, CA, USA) with Sequest HT algorithm and MaxQuant (http://maxquant.org/, version 1.6.0.1). Trypsin/P or Chymotrypsin was selected as the digestive enzyme with two potential missed cleavages. The search included static modification of cysteine carboxymethylation, dynamic modifications of methionine oxidation and serine/threonine/tyrosine phosphorylation.

## Supporting information

Response to Referees Letter

## Acknowledgments

The cryo-EM data were collected at the Center of Cryo-Electron Microscopy, Shanghai Institute of Material Medica. We are also grateful to the staff of the National Center for Protein Science (Shanghai) Electron Microscopy facility for instrument support. We thank the Institutional Technology Service Center of Shanghai Institute of Materia Medica, Chinese Academy of Sciences for technical assistance in mass spectrometry experiments and analysis.

## Author contributions

W.Y. designed the expression constructs, purified the Arr2–NTSR1 complex, prepared the final samples for negative stain and data collection toward the structures, design functional assays, performed disulfide crosslinking, and participated in figure and manuscript preparation. Z.L. performed cryo-EM data collection, processing, map refinement and figure preparation; M. J. performed cryo-EM sample screening, initial reconstruction and figure preparation; Y.Y. performed cell-based functional experiments; P.W.d.W. performed MD simulations and figure preparation; K. P., X.G., Y. H. and X. W. performed construct design and disulfide crosslinking; Y.Z. participated in EM map interpretation and figure preparation; J. G. and H. Z. performed mass spectrometry phosphorylation identification; K. M. and Y. J. participated in experimental design and manuscript editing; X.E.Z. built and refined the structure models, prepared figures and wrote the manuscript; X.Y. designed cryo-EM data collection and processing strategy, performed data processing and map refinement, and participated in manuscript editing; H.E.X. conceived and supervised the project, analyzed the structures, and wrote the manuscript.

## Data and materials availability

Density maps and structure coordinates have been deposited to the Electron Microscopy Database and the Protein Data Bank with accession numbers EMD-20505 and PDB ID 6PWC.

---

**Extended Data Fig. 1.**
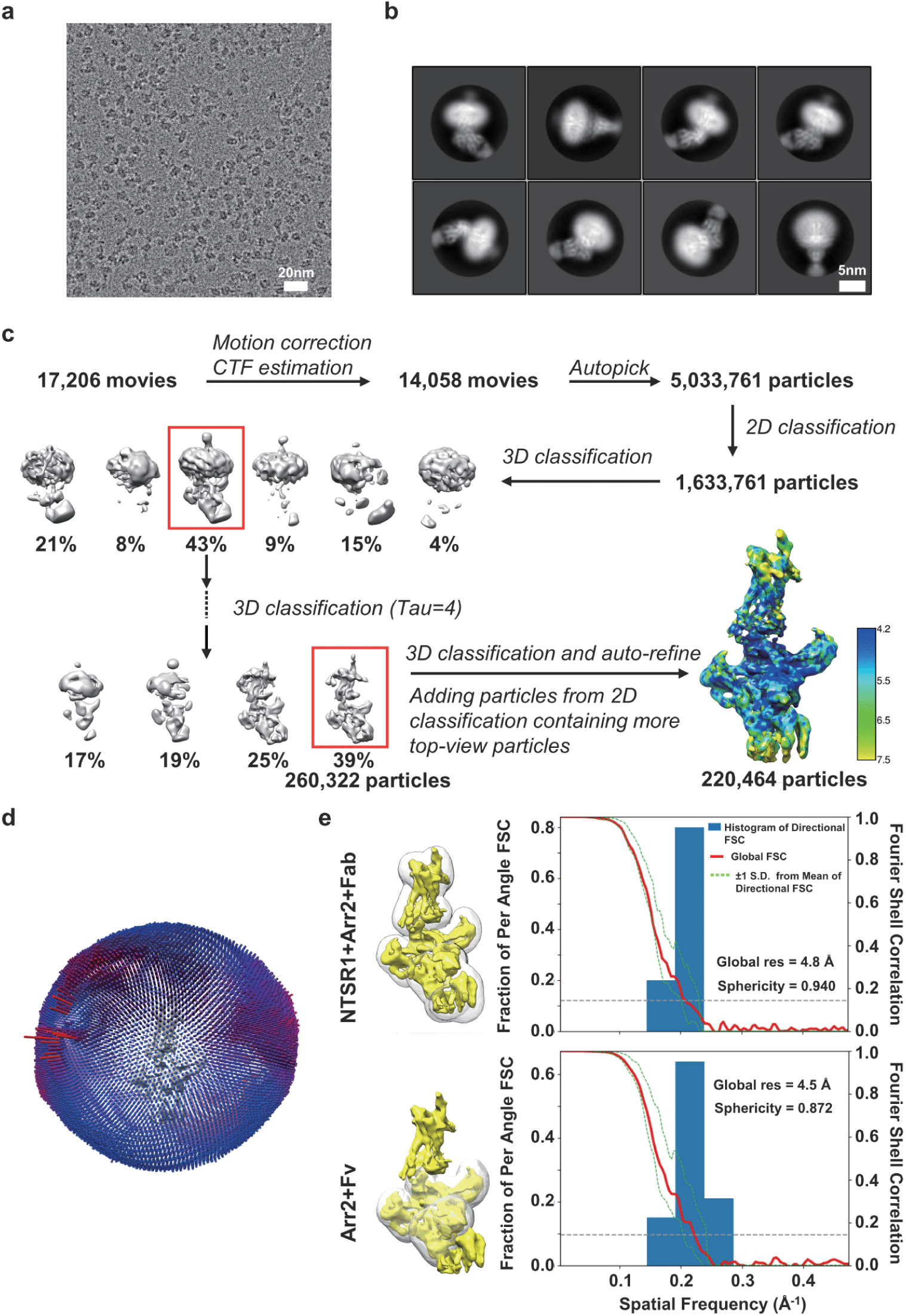
Cryo-EM single-particle analysis of BRIL–NTSR1–Arr2–Fab30 complex. **a**, Representative cryo-EM micrograph of the BRIL–NTSR1–Arr2–Fab30 fusion protein (scale bar, 20 nm). **b**, Representative 2D classes showing different orientations of the BRIL–NTSR1–Arr2–Fab30 fusion protein (scale bar, 5 nm). **c**, Flow chart of data processing with the final reconstruction colored by local resolutions. **d**, Euler angle distribution of particles used in the final reconstruction. **e**, 3D Fourier shell correlation (FSC) analysis. Left, side views of the reconstruction (yellow) superimposed onto the masks (semi-transparent white) produced from the density maps of the whole complex (upper, FSC calculation for the whole complex) and the Arr2-Fv subcomplex (lower, FSC calculation for the subcomplex), respectively. Right, plots of the global half-map FSC (red), the spread of directional resolution values defined by ±1δ from the mean (the area encompassed by dotted green lines, right axis) and the histogram of the directional resolution values evenly sampled over the 3D FSC (blue bars, left axis). The two half maps for the global FSC analysis are produced by 3D auto-refine of Relion 3.0.

**Extended Data Fig. 2.**
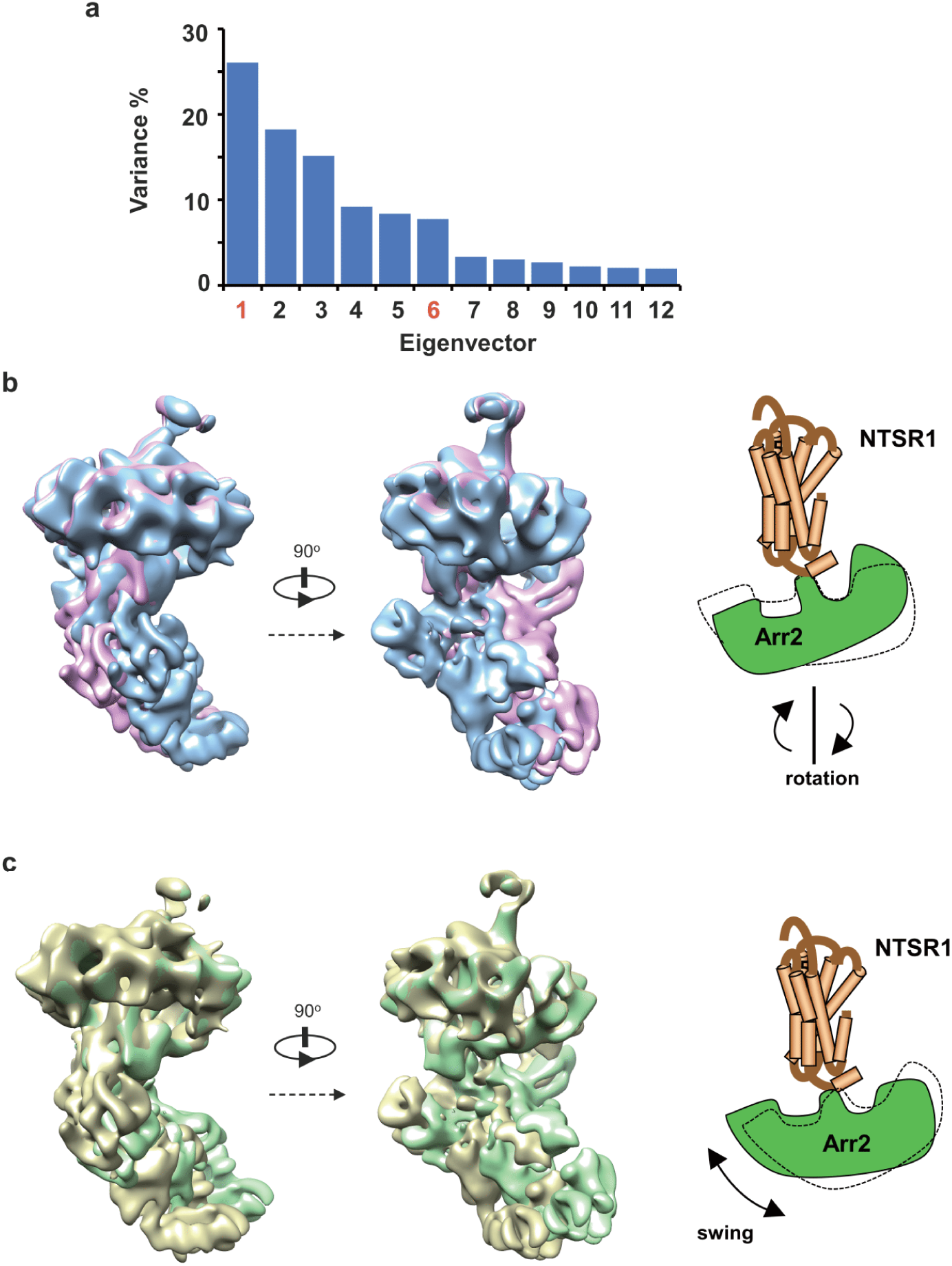
Conformational heterogeneity analysis by multi-body refinement. **a**, The distributional variances of all 12 eigenvectors derived from the particles used for the final reconstruction. The first 6 eigenvectors explain 84.7% of the variance in the dataset. **b**, The two density maps (displayed at 6.5σ) from the 1st and the 10th component in the first eigenvector are aligned by receptor, showing rotational movement of Arr2 with respect to receptor as illustrated by the cartoon. **c**, The two density maps from the 1st and the 10th component in the sixth eigenvector are aligned by receptor, showing swing movement of Arr2 with respect to receptor as illustrated by the cartoon.

**Extended Data Fig. 3.**
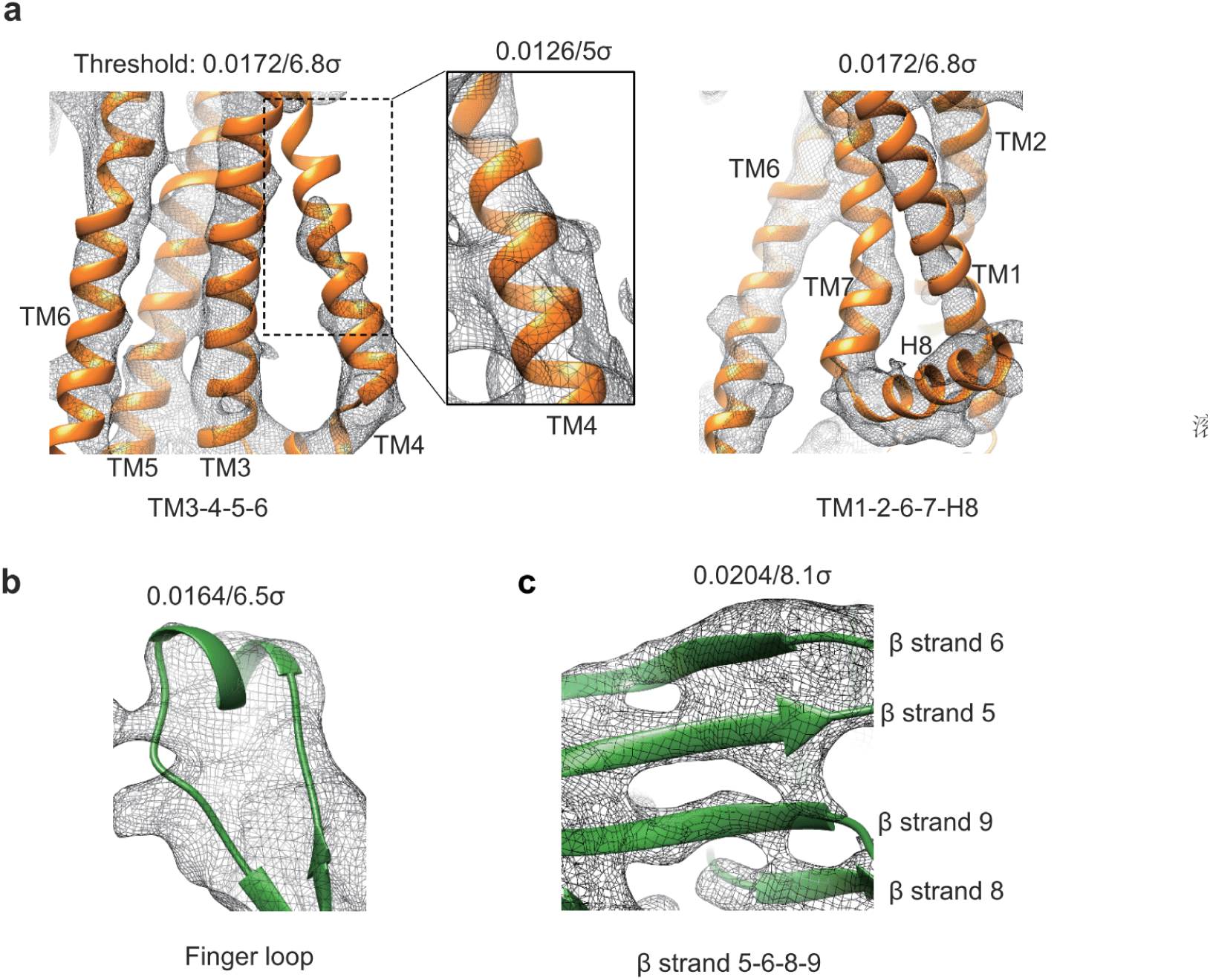
Cryo-EM density map of BRIL–NTSR1–Arr2–Fab30 complex. **a**, Density map (mesh) and the atomic model (ribbon) of NTSR1 with TM helices labeled (the box showing TM4 with smaller contour level to show the continuous density). **b**, Cryo-EM density map of the finger loop of Arr2. **c**, Density map (mesh, low-pass filtered to 4.5 Å with a b-factor of −200 Å^2^) and the atomic model, showing partially separated β-strands (β5, β6, β8 and β9) of Arr2. NTSR1 is colored in brown, and Arr2 finger loop in green. Map contour levels are labeled in the figure panels.

**Extended Data Fig. 4.**
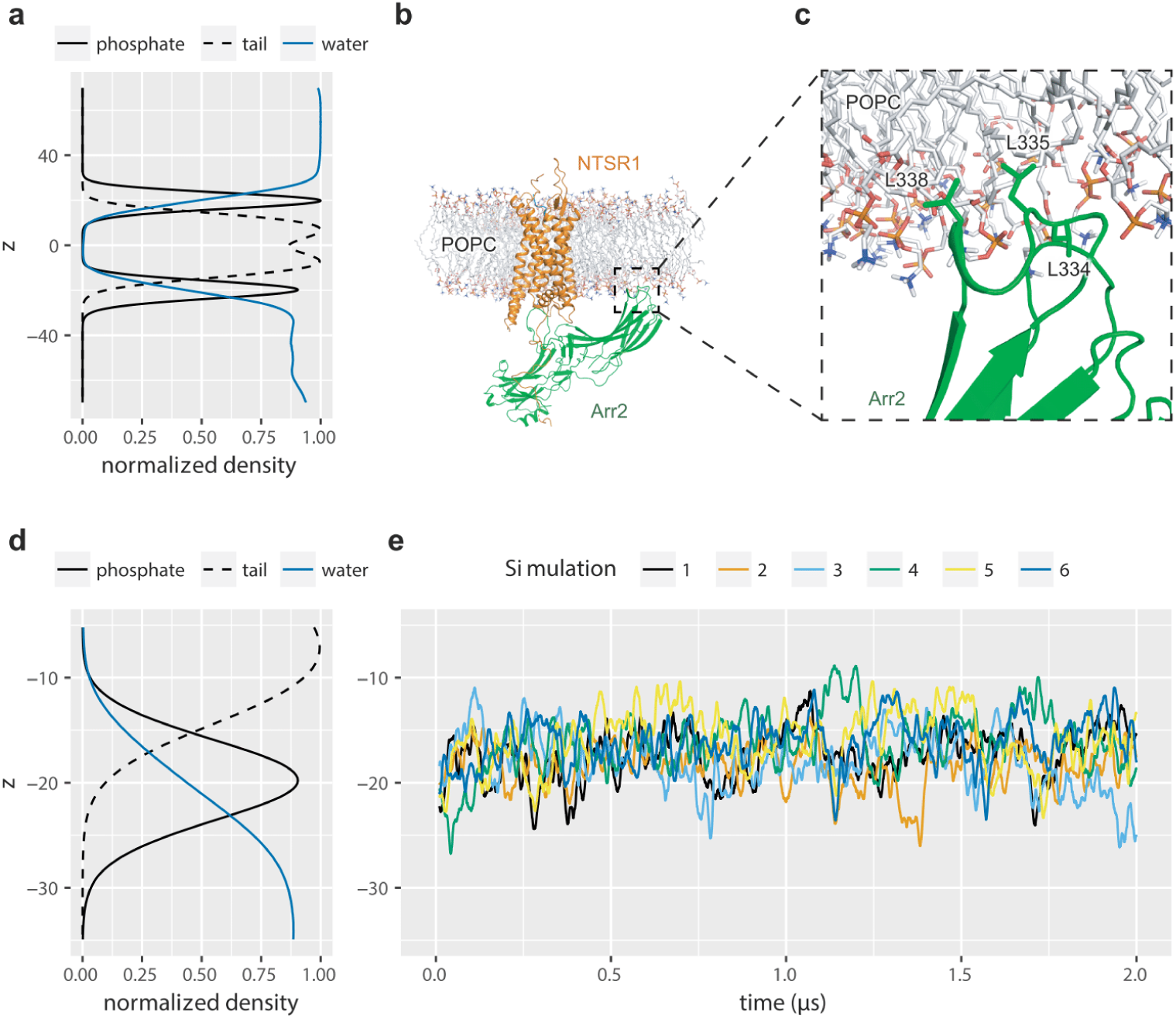
C-edge loop (L334 through L338) of Arr2 associates with the lipid bilayer. **a-b,** Normalized lipid and water solvent density and aligned system snapshot taken from simulation 1. **c,** Three leucine residues (L334, L335 and L338) on the arrestin C-edge loop provide a hydrophobic anchor in lipid bilayer. **d,** Zoomed-in normalized system density near the anchoring C-edge of arrestin. **e,** Average z-axis coordinates of heavy atoms of L334, L335 and L338 shown in (c).

**Extended Data Fig. 5.**
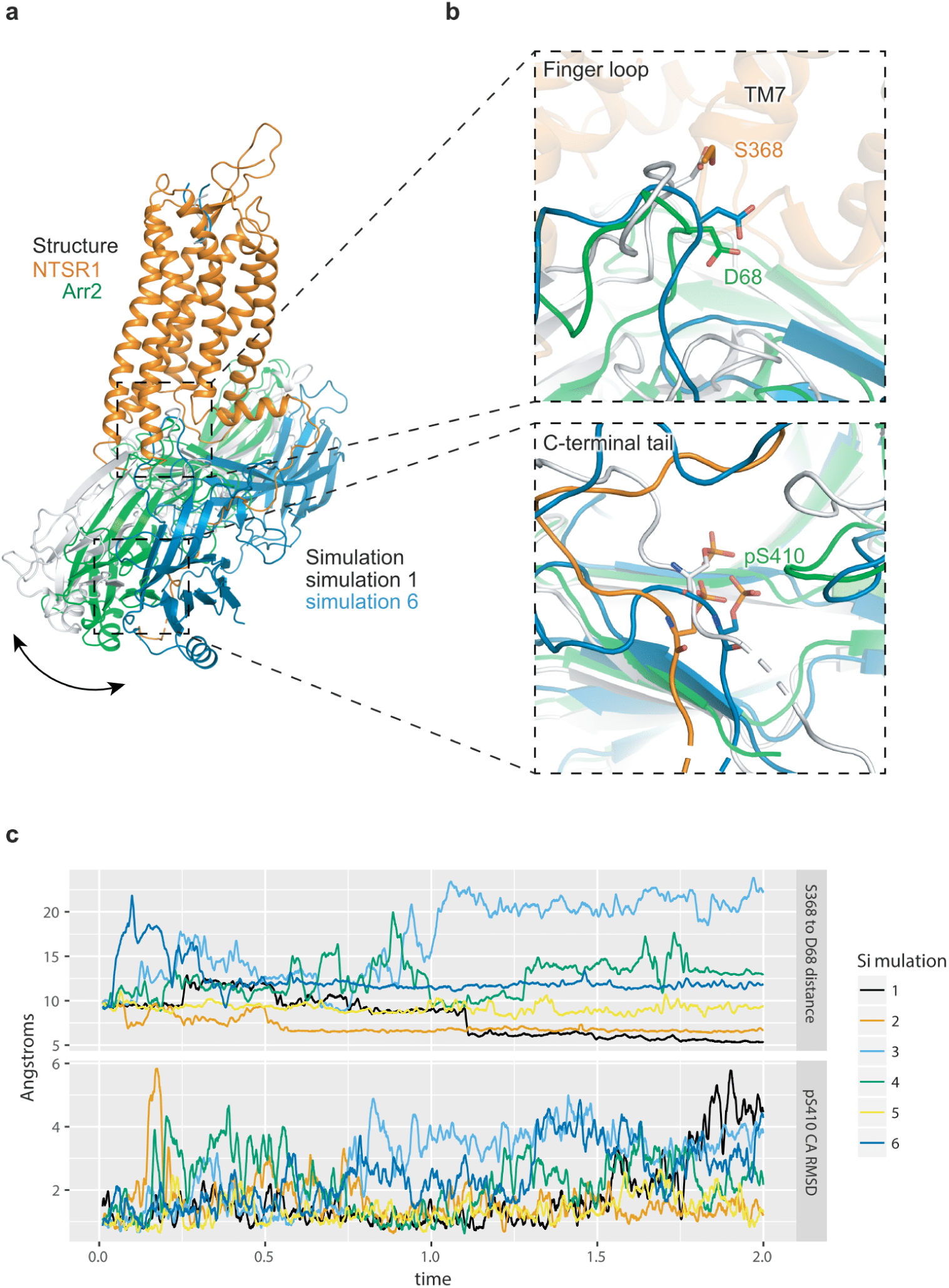
The stability of the interface between Arr2 and NTSR1. **a,** Structural alignment of the cryo-EM Arr2-NTSR1 complex (NTSR1 in brown, Arr2 in green) with two representative simulations (simulations 1 in gray and simulation 6 in blue) terminal poses aligned by the NTSR1 TMD. **b,** Comparison of the terminal positions of finger loop and the receptor C-terminal residue S410 from two simulations displayed in (a). For the S410 comparison, structures were aligned based on the arrestin molecule. **c,** Distance between S368 and D68 (the finger loop interface), and S410 RMSD of six simulations. RMSD calculations were performed relative to the arrestin molecule.

**Extended Data Fig. 6.**
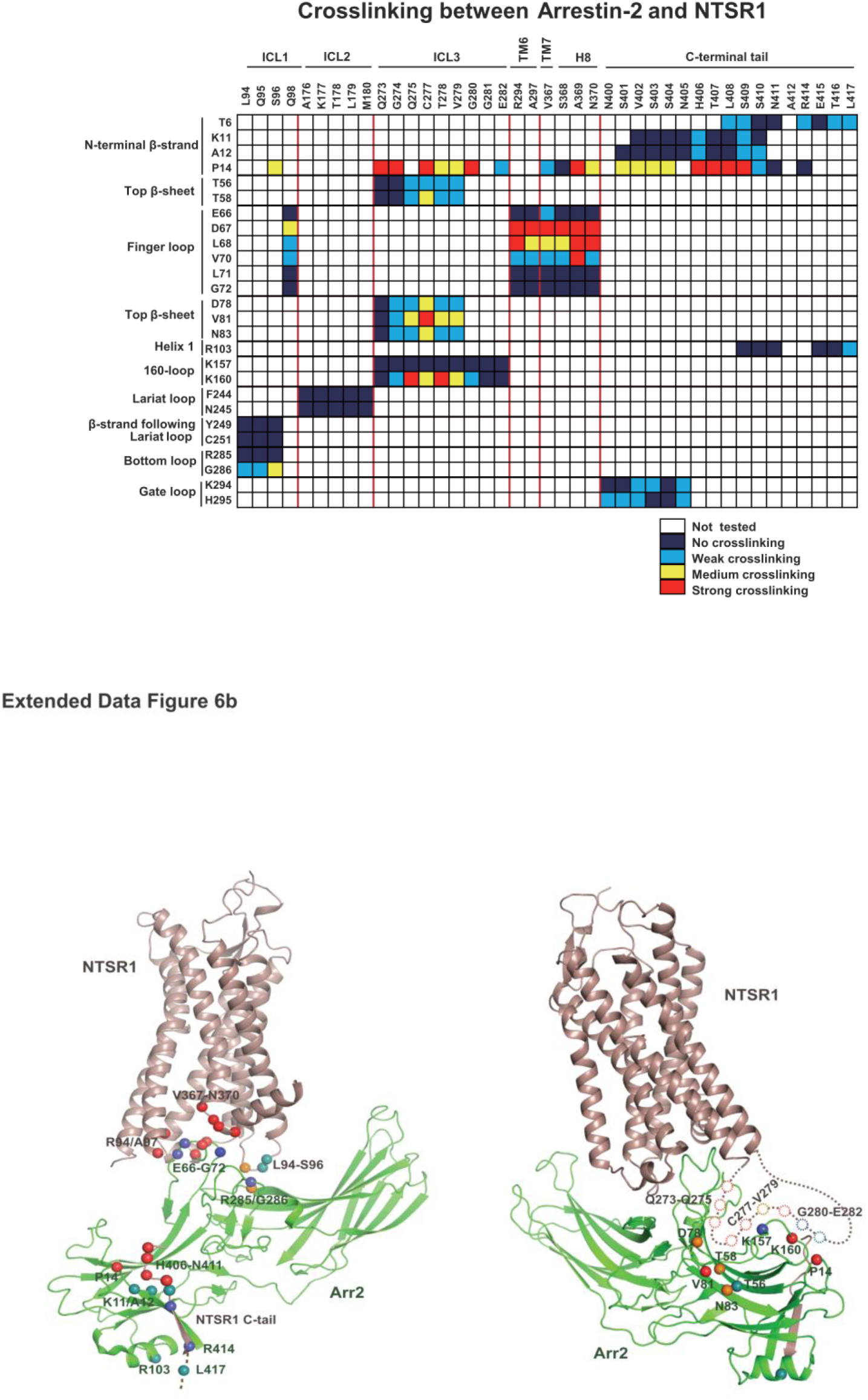
Heat map summary of all Arr2–NTSR1disulfide crosslinking data. **a**, Heat map summary of all Arr2–NTSR1disulfide crosslinking data. **b**, Mapping of the disulfide crosslinking residues at the interface of Arr2-NTSR1 complex. Color code used for the strength of crosslinking signals: strong crosslinking is shown with red (top 75-100%, where 100% mean the strongest crosslinking signal); medium crosslinking, yellow (~50%); weak crosslinking, cyan (~25%); and no obvious crosslinking signal, dark blue (0%, no crosslinking signal). The color for the strongest crosslinking was used when a residue was used for multiple crosslinking pairs.

**Extended Data Fig. 7.**
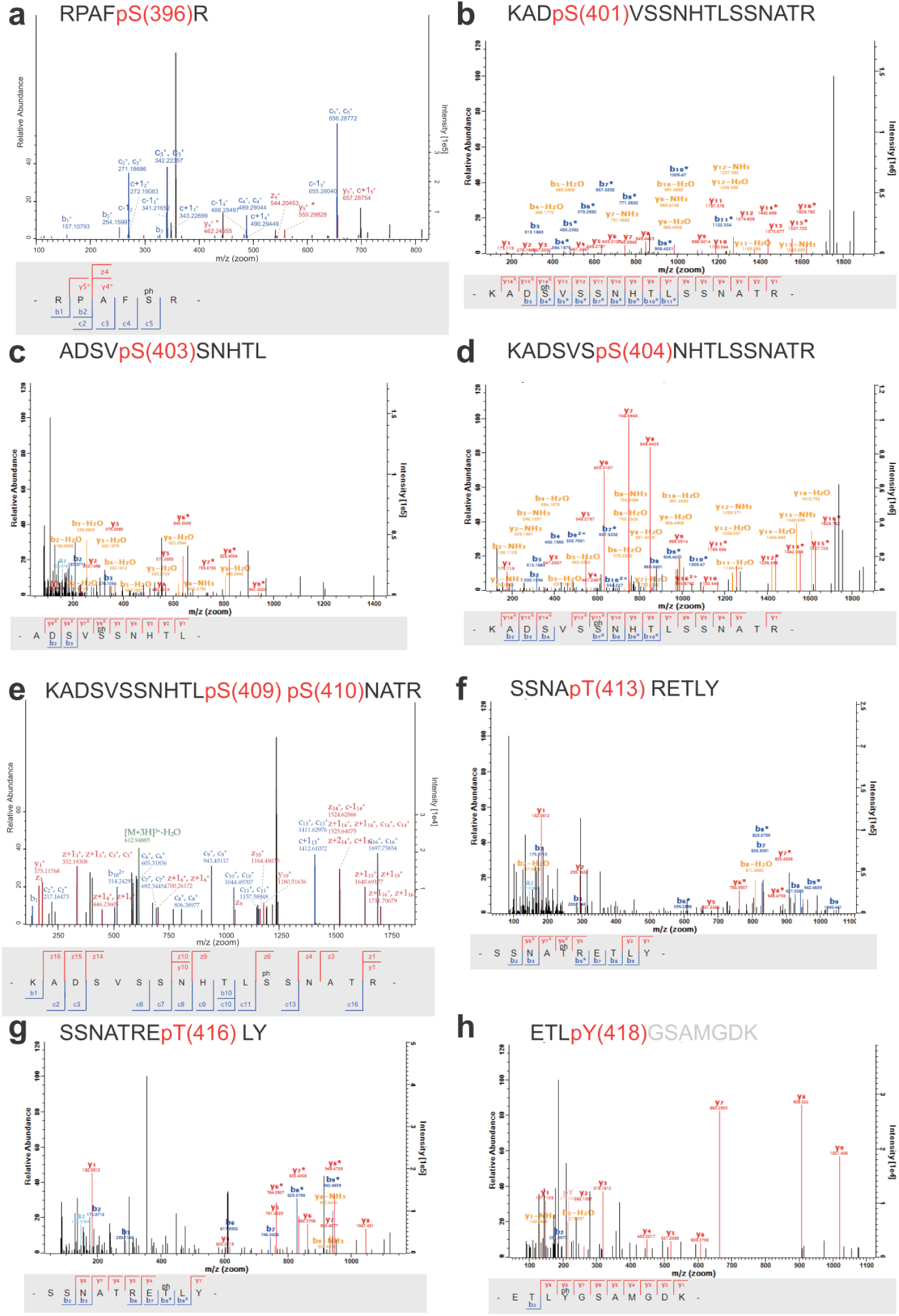
Liquid Chromatography-Mass Spectrometry analysis of BRIL–NTSR1–Arr2–Fab30 fusion protein expressed in SF9. Mass spectra of the BRIL–NTSR1–Arr2–Fab30 fusion protein indicated that the residues S396 (a), S401 (b), S403 (c), S404 (d), S409 or S410 (**e**), T413 (**f**), T416 (**g**), and Y418 (**h**) were phosphorylated. **a** and **e** are ETHCD fragmentation spectrum of the phosphopeptides of corresponding residues. **b, c, d, f, g** and **h** are HCD fragmentation spectra of the phosphopeptides of corresponding residues. ‘ph’ represents a possible phosphate modification. Grey amino acids in (h) represent the additional coding sequence from plasmid.

**Extended Data Fig. 8.**
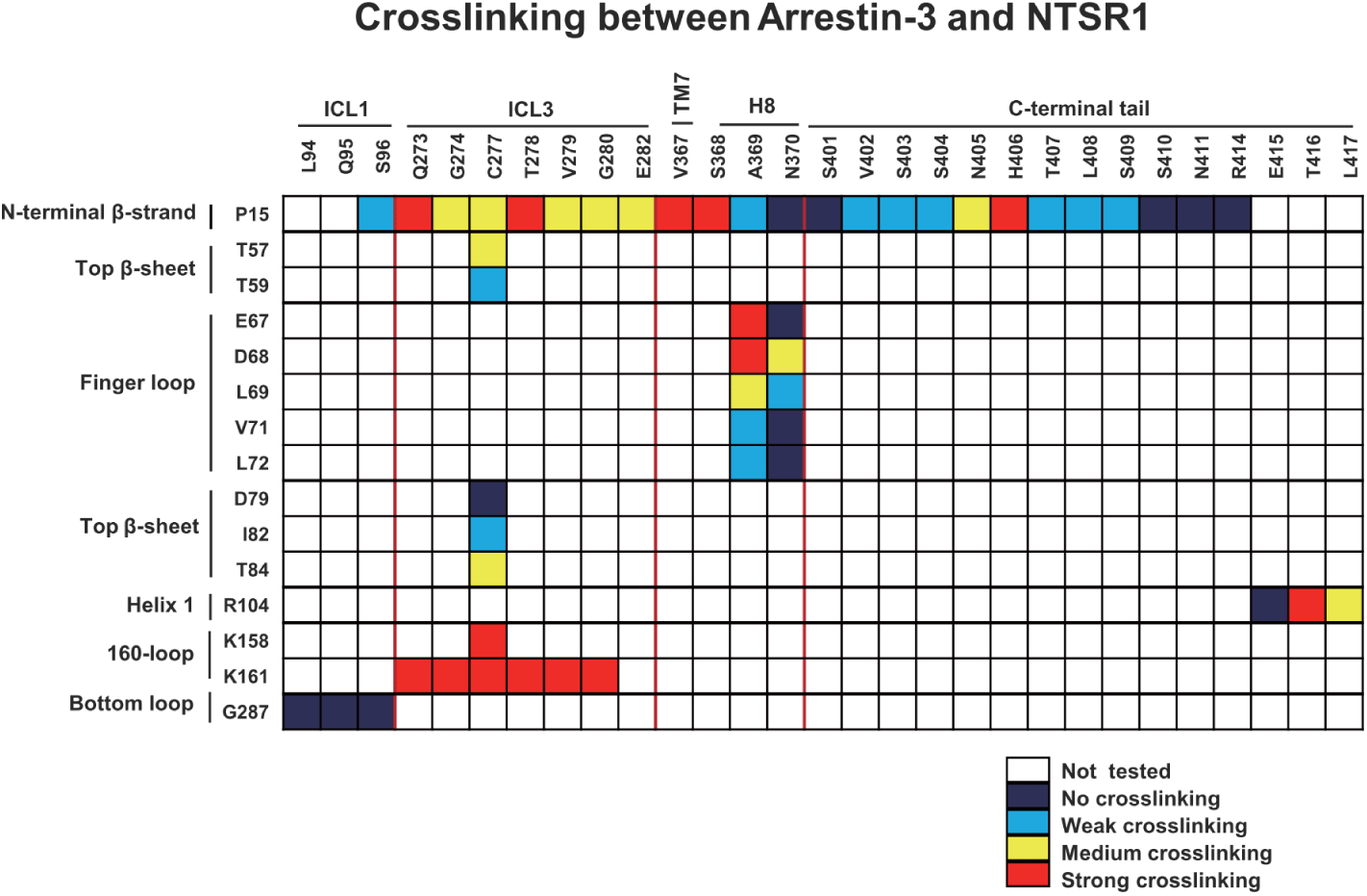
Heat map summary of all Arr3–NTSR1disulfide crosslinking data. Color code used for the strength of crosslinking signals: strong crosslinking is shown with red; medium crosslinking, yellow; weak crosslinking, cyan; and no obvious crosslinking signal, dark blue.

**Extended Data Fig. 9.**
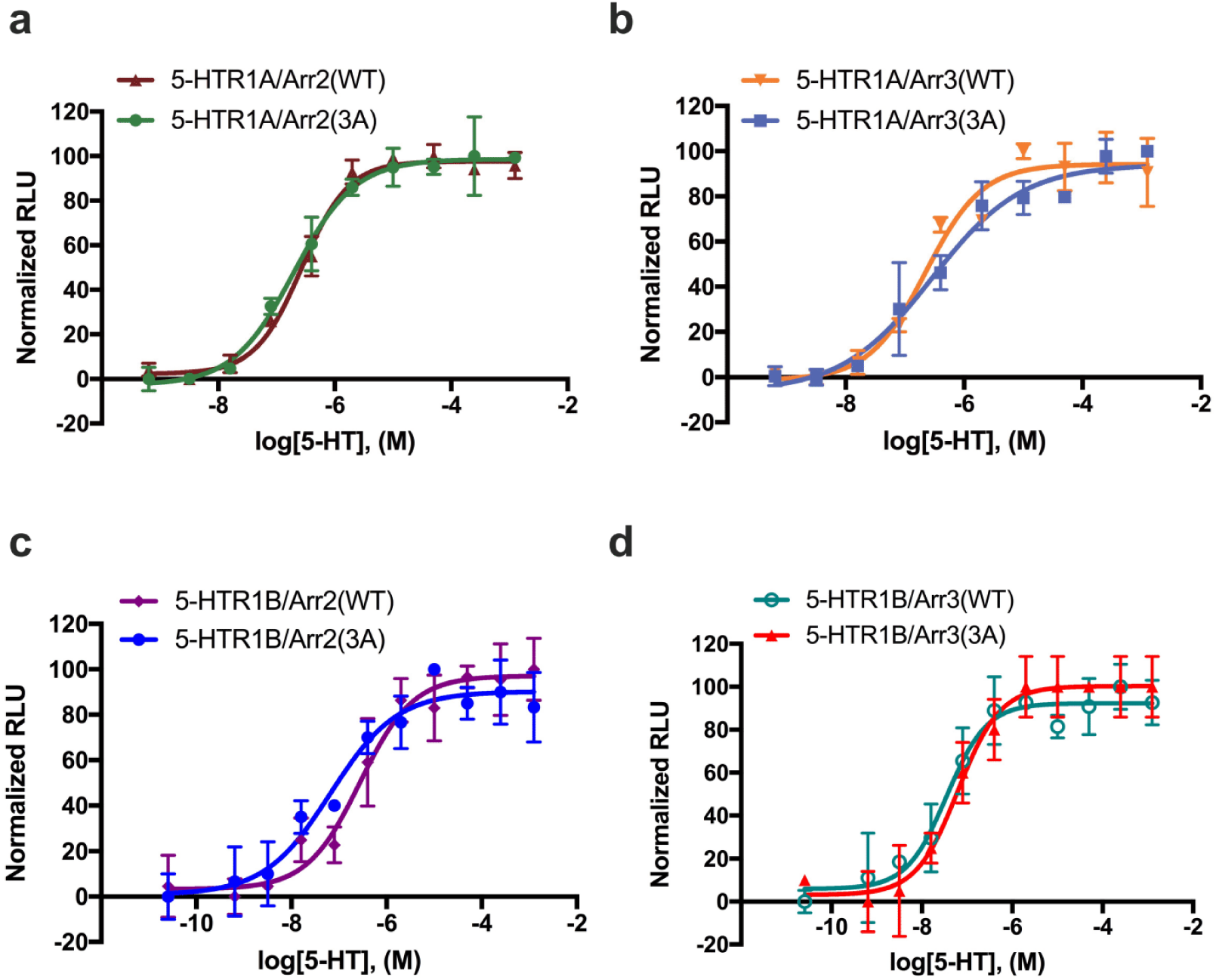
NanoBiT assay to determine β-arrestin recruitment by 5-HTR1A/1B. Concentration–response curves of Arr2 (wild-type and 3A mutant) recruitment by 5HTR1A (**a**), Arr3 (wild-type and 3A mutant) recruitment by 5HTR1A (**b**), Arr2 (wild-type and 3A mutant) recruitment by 5HTR1B (**c**), Arr3 (wild-type and 3A mutant) recruitment by 5HTR1B (**d**). Symbols and error bars represent mean and s.e.m. of indicated independent numbers of experiments(n=3).

**Extended Data Fig. 10.**
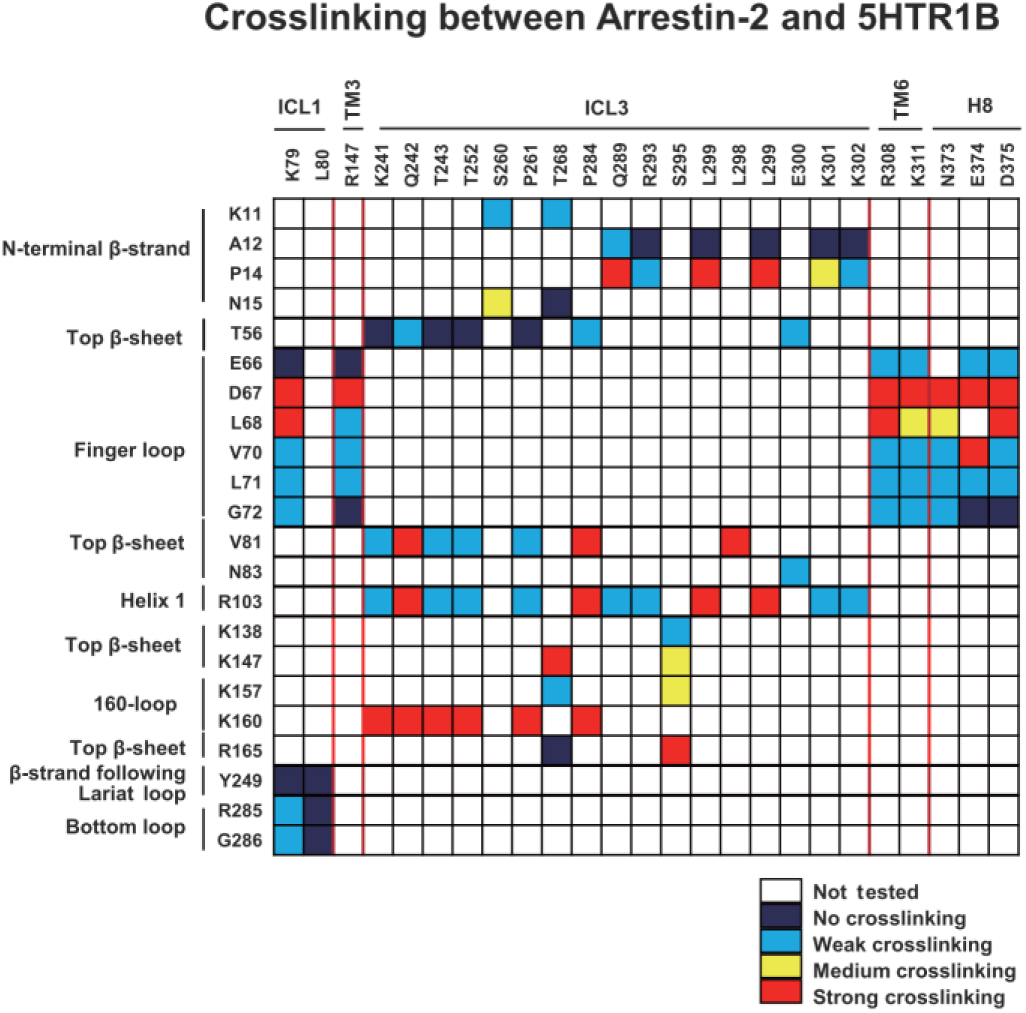
Heat map summary of all Arr2–5-HTR1B disulfide crosslinking data. Color code used for the strength of crosslinking signals: strong crosslinking is shown with red; medium crosslinking, yellow; weak crosslinking, cyan; and no obvious crosslinking signal, dark blue.

**Extended Data Fig. 11.**
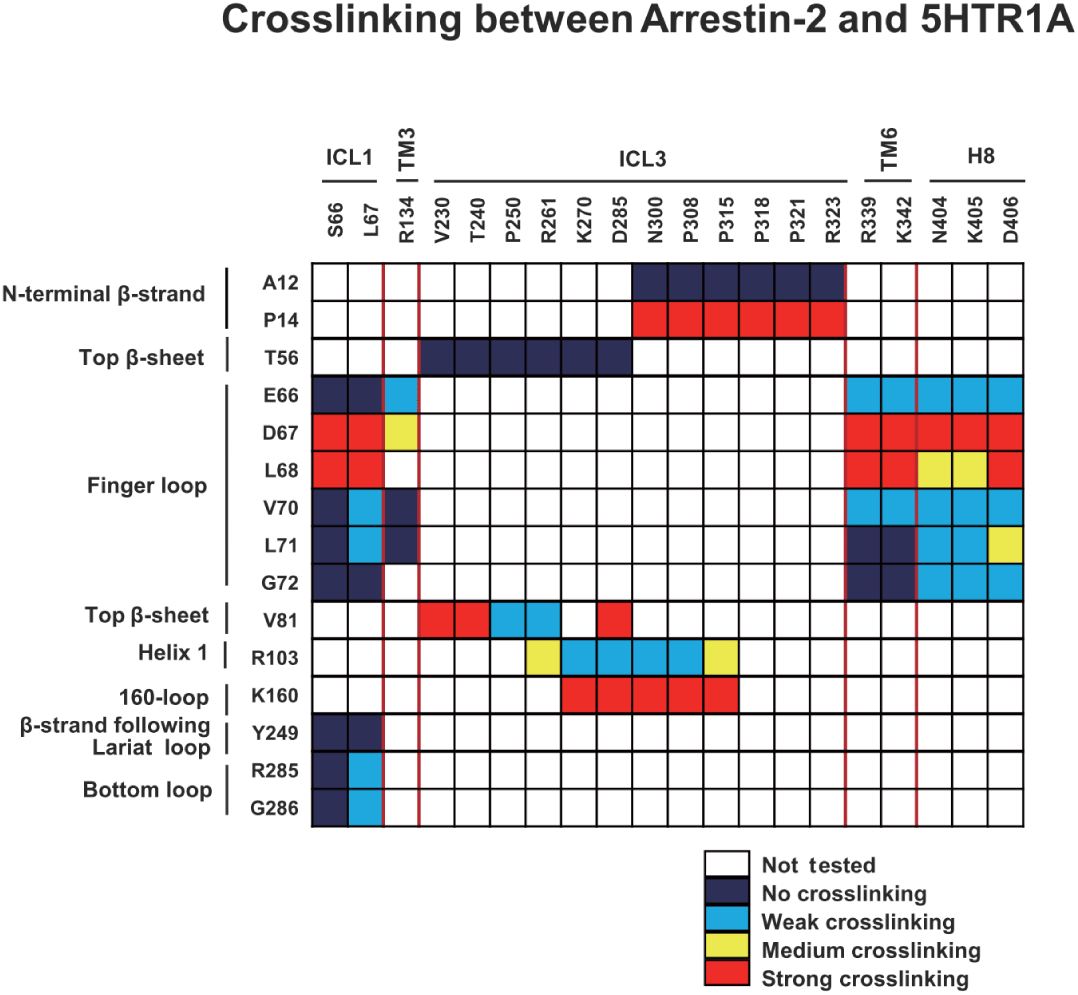
Heat map summary of all Arr2–5-HTR1A disulfide crosslinking data. Color code used for the strength of crosslinking signals: strong crosslinking is shown with red; medium crosslinking, yellow; weak crosslinking, cyan; and no obvious crosslinking signal, dark blue.

**Extended Data Fig. 12.**
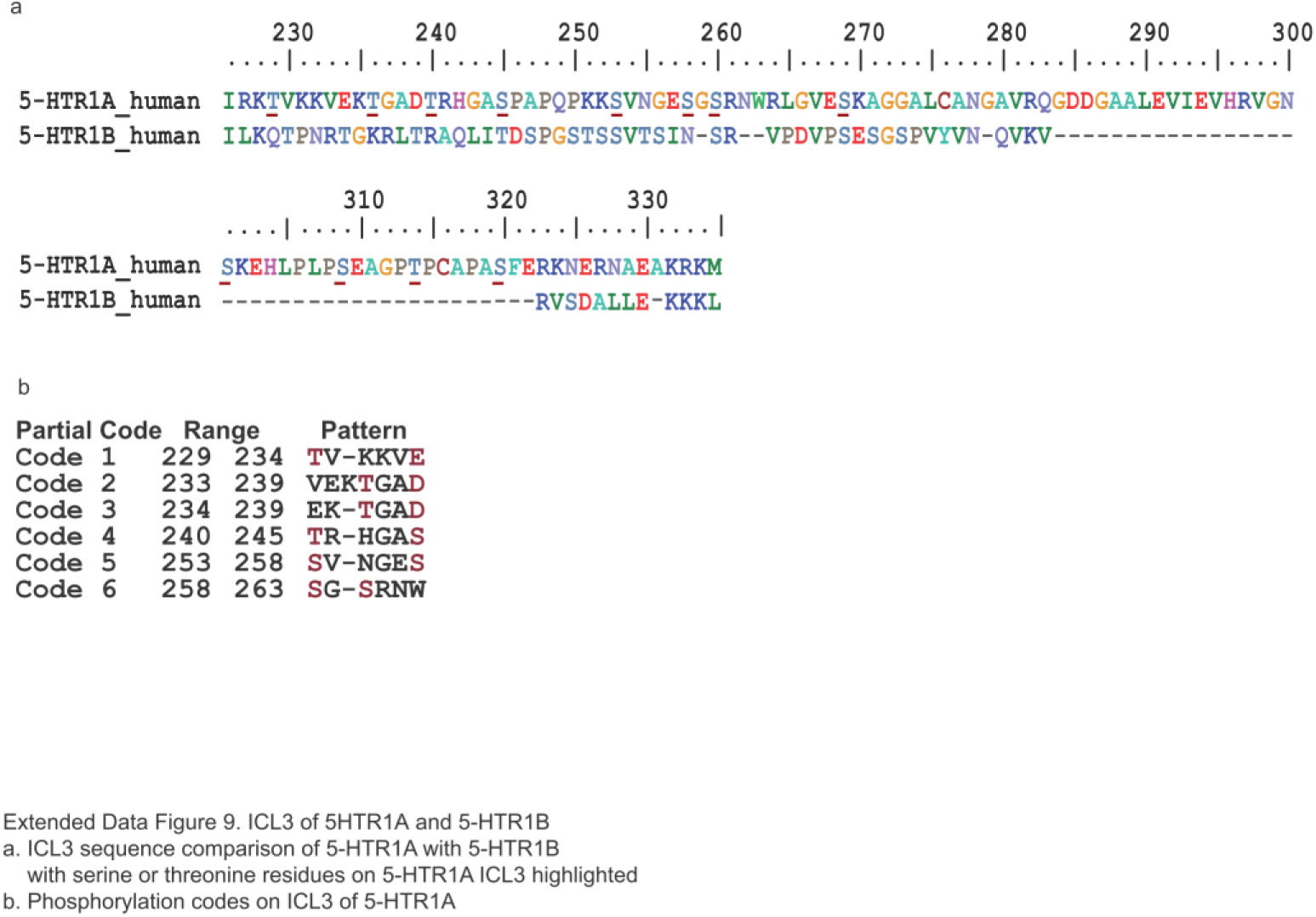
ICL3 of 5HTR1A and 5-HTR1B. **a**, ICL3 sequence comparison of 5-HTR1A with 5-HTR1B with serine or threonine residues on 5-HTR1A ICL3 highlighted. **b**, Phosphorylation codes on ICL3 of 5-HTR1A.

**Extended Data Table 1.**

|  | NTSR1-Arr2-Fab30<br>(EMDB-20505; PDB 6PWC) |
| --- | --- |
| **Data collection and processing** |  |
| Magnification | 48,497 |
| Voltage (kV) | 300 |
| Electron exposure (e<sup>-</sup>/Å<sup>2</sup>) | 68 |
| Defocus range (μm) | -2.0 ~ -3.5 |
| Pixel size (Å) | 1.031 |
| Symmetry imposed | C1 |
| Initial particle images (no.) | 4,960,230 |
| Final particle images (no.) | 220,464 |
| Map resolution (Å) | 4.8 |
| FSC threshold | 0.143 |
| Map resolution range (Å) | 4.5~7.5 |
| **Refinement** |  |
| Initial model used (PDB code) | 4GRV, 4JQI |
| Model resolution (Å) | 4.8 |
| FSC threshold | 0.143 |
| Model resolution range (Å) | 50-4.8 |
| Map sharpening *B* factor (Å<sup>2</sup>) | -250 |
| Model composition |  |
| Non-hydrogen atoms | 5879 |
| Protein residues | 1060 |
| *B* factors (Å<sup>2</sup>) |  |
| Protein | 302 |
| R.M.S. deviations |  |
| Bond lengths (Å) | 0.005 |
| Bond angles (°) | 1.14 |
| Validation |  |
| MolProbity score | 0.98 |
| Clash score | 2.08 |
| Poor rotamers (%) | 0.00 |
| Ramachandran plot |  |
| Favored (%) | 98.8 |
| Allowed (%) | 1.2 |
| Disallowed (%) | 0.00 |

## References

1 Shukla, A. K., Xiao, K. & Lefkowitz, R. J. Emerging paradigms of beta-arrestin-dependent seven transmembrane receptor signaling. Trends in biochemical sciences 36, 457–469, doi:10.1016/j.tibs.2011.06.003 (2011).

2 Ritter, S. L. & Hall, R. A. Fine-tuning of GPCR activity by receptor-interacting proteins. Nature reviews. Molecular cell biology 10, 819–830, doi:10.1038/nrm2803 (2009).

3 Gurevich, V. V. & Gurevich, E. V. Molecular Mechanisms of GPCR Signaling: A Structural Perspective. International journal of molecular sciences 18, doi:10.3390/ijms18122519 (2017).

4 Gurevich, V. V. & Gurevich, E. V. Structural determinants of arrestin functions. Prog Mol Biol Transl Sci 118, 57–92, doi:10.1016/B978-0-12-394440-5.00003-6 (2013).

5 Kang, D. S., Tian, X. & Benovic, J. L. Role of beta-arrestins and arrestin domain-containing proteins in G protein-coupled receptor trafficking. Curr. Opin. Cell Biol. 27, 63–71, doi:10.1016/j.ceb.2013.11.005 (2014).

6 Zhou, X. E., Melcher, K. & Xu, H. E. Understanding the GPCR biased signaling through G protein and arrestin complex structures. Current opinion in structural biology 45, 150–159, doi:10.1016/j.sbi.2017.05.004 (2017).

7 Kang, Y. et al. Crystal structure of rhodopsin bound to arrestin by femtosecond X-ray laser. Nature 523, 561–567, doi:10.1038/nature14656 (2015).

8 Zhou, X. E. et al. X-ray laser diffraction for structure determination of the rhodopsin-arrestin complex. Scientific data 3, 160021, doi:10.1038/sdata.2016.21 (2016).

9 Zhou, X. E. et al. Identification of Phosphorylation Codes for Arrestin Recruitment by G Protein-Coupled Receptors. Cell 170, 457–469 e413, doi:10.1016/j.cell.2017.07.002 (2017).

10 Zhou, X. E., Melcher, K. & Xu, H. E. Structural biology of G protein-coupled receptor signaling complexes. Protein Sci 28, 487–501, doi:10.1002/pro.3526 (2019).

11 Shukla, A. K. et al. Visualization of arrestin recruitment by a G-protein-coupled receptor. Nature 512, 218–222, doi:10.1038/nature13430 (2014).

12 Tanaka, K., Masu, M. & Nakanishi, S. Structure and functional expression of the cloned rat neurotensin receptor. Neuron 4, 847–854 (1990).

13 Kitabgi, P. Targeting neurotensin receptors with agonists and antagonists for therapeutic purposes. Curr Opin Drug Discov Devel 5, 764–776 (2002).

14 White, J. F. et al. Structure of the agonist-bound neurotensin receptor. Nature 490, 508–513, doi:10.1038/nature11558 (2012).

15 Krumm, B. E., White, J. F., Shah, P. & Grisshammer, R. Structural prerequisites for G-protein activation by the neurotensin receptor. Nature communications 6, 7895, doi:10.1038/ncomms8895 (2015).

16 Krumm, B. E. et al. Structure and dynamics of a constitutively active neurotensin receptor. Sci Rep 6, 38564, doi:10.1038/srep38564 (2016).

17 Dixon, A. S. et al. NanoLuc Complementation Reporter Optimized for Accurate Measurement of Protein Interactions in Cells. ACS Chem Biol 11, 400–408, doi:10.1021/acschembio.5b00753 (2016).

18 Gurevich, V. V. The selectivity of visual arrestin for light-activated phosphorhodopsin is controlled by multiple nonredundant mechanisms. The Journal of biological chemistry 273, 15501–15506 (1998).

19 Celver, J., Vishnivetskiy, S. A., Chavkin, C. & Gurevich, V. V. Conservation of the phosphate-sensitive elements in the arrestin family of proteins. The Journal of biological chemistry 277, 9043–9048, doi:10.1074/jbc.M107400200 (2002).

20 Barnea, G. et al. The genetic design of signaling cascades to record receptor activation. Proceedings of the National Academy of Sciences of the United States of America 105, 64–69, doi:10.1073/pnas.0710487105 (2008).

21 Peddibhotla, S. et al. Discovery of ML314, a Brain Penetrant Non-Peptidic beta-Arrestin Biased Agonist of the Neurotensin NTR1 Receptor. ACS Med Chem Lett 4, 846–851, doi:10.1021/ml400176n (2013).

22 Barak, L. S. et al. ML314: A Biased Neurotensin Receptor Ligand for Methamphetamine Abuse. ACS Chem Biol 11, 1880–1890, doi:10.1021/acschembio.6b00291 (2016).

23 Shukla, A. K. et al. Structure of active beta-arrestin-1 bound to a G-protein-coupled receptor phosphopeptide. Nature 497, 137–141, doi:10.1038/nature12120 (2013).

24 Alexandrov, A. I., Mileni, M., Chien, E. Y. T., Hanson, M. A. & Stevens, R. C. Microscale fluorescent thermal stability assay for membrane proteins. Structure 16, 351–359 (2008).

25 Wang, R. Y. et al. Automated structure refinement of macromolecular assemblies from cryo-EM maps using Rosetta. Elife 5, doi:10.7554/eLife.17219 (2016).

26 Lally, C. C., Bauer, B., Selent, J. & Sommer, M. E. C-edge loops of arrestin function as a membrane anchor. Nature communications 8, 14258, doi:10.1038/ncomms14258 (2017).

27 Sommer, M. E., Hofmann, K. P. & Heck, M. Distinct loops in arrestin differentially regulate ligand binding within the GPCR opsin. Nature communications 3, 995, doi:10.1038/ncomms2000 (2012).

28 Janoshazi, A. et al. Modified receptor internalization upon coexpression of 5-HT1B receptor and 5-HT2B receptors. Molecular pharmacology 71, 1463–1474, doi:10.1124/mol.106.032656 (2007).

29 Wang, C. et al. Structural basis for molecular recognition at serotonin receptors. Science 340, 610–614, doi:10.1126/science.1232807 (2013).

30 Zheng, S. Q. et al. MotionCor2: anisotropic correction of beam-induced motion for improved cryo-electron microscopy. Nat Methods 14, 331–332, doi:10.1038/nmeth.4193 (2017).

31 Zhang, K. Gctf: Real-time CTF determination and correction. J Struct Biol 193, 1–12, doi:10.1016/j.jsb.2015.11.003 (2016).

32 Zivanov, J. et al. New tools for automated high-resolution cryo-EM structure determination in RELION-3. Elife 7, doi:10.7554/eLife.42166 (2018).

33 Kucukelbir, A., Sigworth, F. J. & Tagare, H. D. Quantifying the local resolution of cryo-EM density maps. Nat Methods 11, 63–65, doi:10.1038/nmeth.2727 (2014).

34 Pettersen, E. F. et al. UCSF chimera - A visualization system for exploratory research and analysis. J Comput Chem 25, 1605–1612 (2004).

35 Emsley, P. & Cowtan, K. Coot: model-building tools for molecular graphics. Acta Crystallogr D 60, 2126–2132, doi:Doi 10.1107/S0907444904019158 (2004).

36 Adams, P. D. et al. PHENIX: a comprehensive Python-based system for macromolecular structure solution. Acta crystallographica. Section D, Biological crystallography 66, 213–221, doi:10.1107/S0907444909052925 (2010).

37 DiMaio, F. et al. Improved low-resolution crystallographic refinement with Phenix and Rosetta. Nat Methods 10, 1102–1104, doi:10.1038/nmeth.2648 (2013).

38 Chen, V. B. et al. MolProbity: all-atom structure validation for macromolecular crystallography. Acta crystallographica. Section D, Biological crystallography 66, 12–21, doi:10.1107/S0907444909042073 (2010).

39 Huang, J. et al. CHARMM36m: an improved force field for folded and intrinsically disordered proteins. Nat Methods 14, 71–73, doi:10.1038/nmeth.4067 (2017).

40 D.A. Case, I. Y. B.-S., S.R. Brozell, D.S. Cerutti, T.E. Cheatham, III, V.W.D. Cruzeiro, T.A. Darden,, R.E. Duke, D. G., M.K. Gilson, H. Gohlke, A.W. Goetz, D. Greene, R Harris, N. Homeyer, Y. Huang,, S. Izadi, A. K., T. Kurtzman, T.S. Lee, S. LeGrand, P. Li, C. Lin, J. Liu, T. Luchko, R. Luo, D.J., Mermelstein, K. M. M., Y. Miao, G. Monard, C. Nguyen, H. Nguyen, I. Omelyan, A. Onufriev, F. Pan, R.& Qi, D. R. R., A. Roitberg, C. Sagui, S. Schott-Verdugo, J. Shen, C.L. Simmerling, J. Smith, R. SalomonFerrer, J. Swails, R.C. Walker, J. Wang, H. Wei, R.M. Wolf, X. Wu, L. Xiao, D.M. York and P.A. Kollman. AMBER 2018. Unviersity of California, San Francisco (2018).

41 Lomize, M. A., Lomize, A. L., Pogozheva, I. D. & Mosberg, H. I. OPM: orientations of proteins in membranes database. Bioinformatics 22, 623–625, doi:10.1093/bioinformatics/btk023 (2006).

42 Eswar, N. et al. Comparative protein structure modeling using Modeller. Curr Protoc Bioinformatics Chapter 5, Unit-5 6, doi:10.1002/0471250953.bi0506s15 (2006).

43 McGibbon, R. T. et al. MDTraj: A Modern Open Library for the Analysis of Molecular Dynamics Trajectories. Biophys J 109, 1528–1532, doi:10.1016/j.bpj.2015.08.015 (2015).

44 Humphrey, W., Dalke, A. & Schulten, K. VMD: visual molecular dynamics. J. Mol. Graph. 14, 33–38, 27–38 (1996).

45 Roe, D. R. & Cheatham, T. E., 3rd. PTRAJ and CPPTRAJ: Software for Processing and Analysis of Molecular Dynamics Trajectory Data. J Chem Theory Comput 9, 3084–3095, doi:10.1021/ct400341p (2013).

