## Supplementary material for "A complex structure of arrestin-2 bound to a G protein-coupled receptor": Response to Referees Letter

We want to thank the three reviewers for the tremendous efforts in evaluating our manuscript and for the comments on the importance and biological significance of arrestin binding to non-visual GPCRs. We want to point out that we have been communicating with Kobilka and Skiniotis for the past several months and planning the co-submission of our three papers together. Our works were completed a few weeks before theirs and we have waited for them for several weeks. Eventually, they agreed that we can submit our paper on July 28th, a couple weeks before theirs, even though our paper was completed in Mid-July.

As positively commented by all three reviewers, the most striking observation in the arrestin2 (Arr2)-NTSR1 complex structure is the ~90-degree rotation of Arr2 from the visual arrestin-rhodopsin complex. The overall orientation of NTSR1, ARR2 and Fab30 can be unambiguously placed in our complex. After submission of our paper, Kobilka/Skiniotis and our group exchanged our manuscripts and PDB coordinates, which confirmed that our group has independently reached the similar conclusions, despite of using very different engineering approaches. Kobilka/Skiniotis used PIP2 as well as a truncated form of arrestin with extensive chemical and photo-activated crosslinking to stabilize the complex while our group used the fusion of the receptor with the full-length arrestin and antibody Fab30. In addition, we have further confirmed our structure with extensive site-specific disulfide crosslinking in cells using non-fused proteins in the absence of Fab30. Thus, Kobilka/Skiniotis have determined essentially the same structure using very different engineering approaches, which provides another level of validation. Since these papers complement and validate each other for an extremely challenging structure, the sum of both papers is stronger than either one on its own and they would therefore benefit the readers of Nature from being published together back-to-back.

In the following sections, we will provide point-by-point responses to the comments by the reviewers. The reviewer's comments are in **black** and our responses are in **blue**.

**Response to the comments by Reviewer #1:**
Referee #1 (Remarks to the Author):
Yin et al. present the cryo-EM structure of NTSR1 in complex with arrestin2. To achieve this, they used NTSR1 fused with human arrestin2("3A" a constitutively active mutant), in turn fused to Fab30 (antibody fragment that binds to arrestin2-GPCR tail complex). Cysteine mutants were introduced into arrestin2 to crosslink with IC3 of receptor (V81C) and with Fab30 (A279C), although unclear if both formed in the final product. Used digitonin micelles to solubilize the complex, which is not very plasmid membrane-like. Co-expressed in insect cells with GRK5 to achieve phosphorylation. The final maps, which used a very small fraction of the total particles collected, where of moderate resolution and thus do not reveal features such as side chains nor bound peptide hormone. They confirm their configuration with disulfide mapping, although, like any other crosslinking approach, these results are difficult to interpret because, among other factors, the positions that are the "hottest" often tend to be the most dynamic, as evidenced by positions that hit multiple other sites.
>> The interaction of arrestin to non-visual GPCRs is very transient and dynamic. For the past eight years, our group has devoted exhaustive efforts in testing various methods to stabilize an arrestin/non-visual GPCR complex for structural studies. We eventually succeeded using the fusion construct of Arr2 with Fab30 and NTSR1, coincidently the same receptor used by the Kobilka/Skiniotis groups. The similar structures reached independently by both groups provide additional support for our approaches.

We have also screened extensive detergent conditions and found digitonin and LMNG/GDN mix are the two best conditions for cryo-EM studies. The digitonin micelle is bigger than the LMNG/GDN micelle (thus, digitonin micelles mimic the bilayer of the lipid membrane better than LMNG/GDN micelles) and digitonin is one of the mildest detergents for membrane proteins. In addition, digitonin is one of the most used detergents for membrane protein structure determination by cryo-EM in recent years (https://www.anatrace.com/Landing/2018/Cryo-EM-Update-Oct18). Thus, we feel that digitonin is the appropriate detergent for structure determination of the arrestin-NTSR1 complex.

We agree with the reviewer that crosslinking can occur between multiple positions when they are located in flexible and highly dynamic regions. However, site-specific disulfide crosslinking requires much more specific positions of crosslinking residue pairs and side chain conformation than other types of chemical crosslinking. Thus, in many cases, disulfide crosslinking data can help to identify key interface residues by evaluating the strength of the crosslinking signal.

In addition, disulfide crosslinking experiments were performed in-cell using non-fused receptor and arrestin, which can reveal a complex interface in the more natural condition of a native membrane in cells. Thus, disulfide crosslinking approaches have been widely used for interface mapping and crystal/cryo-EM structure validation. We have used the in-cell disulfide crosslinking technique to validate the complex assembly of the visual arrestin-rhodopsin complex, which results have withstood the test of time since the publication of our original paper in 2015.

The major and most striking result is that although arrestin uses the same structural moiety to bind NTSR1 as it does rhodopsin, the only other published GPCR-arrestin complex, it does so in a manner that results in a ~90˚ rotation of the arrestin relative to the receptor. The authors propose based on cysteine cross linking mapping that other class A receptors will adopt the same configuration and propose it to be a general model for arrestin 2 interactions. The C-lobe of arrestin is situated to interact with the membrane as it does in the rhodopsin-arrestin1 complex.
>>We thank the reviewer for recognizing the two most important results of our paper: 1) a ~90˚ rotation of the arrestin relative to the receptor from the visual arrestin-rhodopsin complex; and 2) The C-lobe of arrestin is situated to interact with the membrane as it does in the rhodopsin-arrestin1 complex. The reviewer's comments have helped us highlight these two central points in our revised manuscript.

Although this is a fine technical achievement and a surprising result, enthusiasm is diminished by the following issues:
1) The map is of poor quality, although adequate for positioning the GPCR, the arrestin, and Fab30. Importantly, the receptor C-tail density is poorly defined and the density for the C-tail portrayed in Fig. 5b is highly misleading and does not correlate well with the provided EM map. Despite what is claimed in the paper, there is no convincing density for phosphorylation at modeled residues Ser409 or Ser410. Crosslinking data seems to indicate a large amount of heterogeneity in the C-tail interaction regardless.
>> We agree with the reviewer's comments that our structure at 4.9Å resolution is adequate for positioning the GPCR, the arrestin, and Fab30, but is insufficient for placing side chains in the structure and phosphorylation residues. As such, we have removed all side chains in our figure illustrations.

We also understand the reviewer's concern about the density of the receptor C-tail. While we see the overall density of the receptor C-tail that binds to the first beta-strand of arrestin, the density for the phosphate groups in the C-tail is unconvincing. Thus, in the revised manuscript, we only present a C-tail region with all side chains truncated (new Fig 5a). We labeled C-tail residues that were validated by disulfide crosslinking to interact with arrestin N-domain (New Fig. 5c). The figure panels and the text about phosphorylation codes are deleted in the revised version.

2) There is no experimental evidence to justify many of the highly detailed structure panels in Figures 3-5 because there is no side chain density in the map at current resolution and interpretation of backbone path is questionable in some important regions like the finger loop.
>> We agree with the reviewer's comment that the density map does not provide highly detailed information due to the limited resolution. In the original manuscript, we showed these side chains to indicate the positions of the crosslinking residues, which we regrettably did not state clearly in the text or in the figure legends. To avoid such confusion, we omitted all side chain-stick presentation of the residues in the structural panels of Figures 3-5 in the revised manuscript. We show all the residues on the backbone trace by small spheres to indicate the positions of the interface residues on the receptor-arrestin interface patches that were supported by disulfide crosslinking data.

During the revision of this manuscript, we went through the whole model and further refined and rebuilt the model against the density map, particularly the interface regions, which fit better into the density. While the side chains of the finger loop are truncated, the density for the backbone trace of the finger loop (E66 through L71) and the flanking beta-strands is clear at contour level of 0.015 (it is the contour level we used for the whole map, which is 5.55 sigma).

3) The discussion of potential phosphocodes should be eliminated because it is distracting from the most important results and the data doesn't really speak to the hypothesis because it is not known what phosphosites are fully occupied or bound to arrestin in the complex, making a discussion of phosphocodes irrelevant. Furthermore, it is unclear if they know just how phosphorylated their fusion protein is (which is co-expressed with GRK5 in Sf9 cells) and we don't know if this fusion is phosphorylated the same way as the native receptor by GRK5 (seems unlikely) or whether all proteins in the sample are phosphorylated in the same manner (even hard to know if all Sf9 cells got both fusion protein and GRK5 viruses). Moreover, the MS/MS data in Extended data Figure 6, although providing evidence for phosphorylation at the indicated Ser/Thr positions, does not and cannot indicate how many of these sites in each protein are simultaneously phosphorylated or the relative amounts of phosphate at each site.
>>We are grateful for this comment of the reviewer, which has help us tremendously improve the manuscript by making our key points much clearer. We completely agree with the reviewer that the potential phosphorylation codes are not relevant to the most important points of this paper. Although we see density at positions of 409 and 413, we understand that these pieces of density do not convincingly represent phosphorylation of these two receptor C-tail positions because of the limited resolution. Therefore, we removed the relevant text and figure panels about phosphorylation of the C-terminal tail of the receptor from the paper.

4) An interesting point brought up by the authors is the possibility that this unusual configuration is how

receptors with phosphorylated IC3 loops and short C-tails interact with arrestins. To this end they studied the HTR1A and B receptor complexes, which have phosphorylated IC3 loops and short C-tails, via disulfide mapping. However, inspection of the rhodopsin-arrestin1 structures suggest that both canonical or NTSR1-like configurations of arrestin are reasonably compatible with much of the disulfide crosslinking results (e.g. crosslink with V81C in panel 6e). These experiments are somewhat flawed because HEK cells do not express GRK5 (Violin et al. 2006). How many IC3 loop sites are getting phosphorylated in the same receptor? It matters because this information would constrain the IC3 loop position depending on where they are located relative to the polar core of arrestin. Thus the hypothesis that this configuration is applicable to other GPCRs seems a stretch.
>> We agree with the reviewer on that many crosslinking data in ICL3 can be reasonably compatible with both orientations of arrestin in the rhodopsin or NTSR1 complex, including L298C-V81C pairs. However, the crosslinking of ICL3 residue S260C of 5-HTR1B with arrestin residue K11 and N15, as well as the crosslinking of ICL1 residue K79C of 5-HTR1B with arrestin residues R285C or G286C are not compatible with the visual arrestin-rhodopsin orientation (Rebuttal Fig.1 in next page, 5HT1A model is based on our unpublished 5HT1A-G protein complex structure). Receptor TM5 is at the back side of arrestin N-domain in the rhodopsin orientation (Rebuttal Fig. 1b&d). It is difficult for ICL3, including S260 and S268, to go over to the front side of arrestin for crosslinking. In contrast, these crosslinking data are compatible with the Arr2-NTSR1 orientation, thus supporting our model of Arr2 coupling by 5HT1B ICL3 (Fig. 6 and Rebuttal Fig. 1a&c).

The crosslinking between ICL1 residue K79C of 5-HTR1B and R285C and G286C of arrestin2(Fig. 6a&b), can only happen in the arrestin2-NTSR1 orientation because in arrestin2-NTSR1 complex structure, the ICL1 of the receptor is positioned in arrestin's top cleft between the top β-sheets of the N- and C-domains of the arrestin where R285 and G286 are located, while the ICL2 of rhodopsin is positioned in this cleft in the visual arrestin-rhodopsin complex structure.

In addition, we also performed distance analysis of crosslinking residue pairs based on the Arr2-NTSR1 orientation and Arr1-Rhodopsin orientation for the crosslinking residues pairs of Arr2 with NTSR1, 5HT1A/1B, as showed in the Rebuttal Table 1-3 attached below. From these distance comparisons, the distances of crosslinking residue pairs are reasonable for disulfide bond formation in the Arr2-NTSR1 orientation and they are much shorter than the corresponding distances in the Arr1-rhodopsin orientation. The distance analyses clearly indicate that the Arr2 orientation in the NTSR1 complex is the only possible model for ICL3 interaction with the N-domain of Arr2, whereas the visual arrestin orientation in the rhodopsin complex is incompatible with the crosslinking data of a number of residue pairs, which are now clearly indicated in the revised manuscript.

On the other hand, we disagree with the comment of "These experiments are somewhat flawed because HEK cells do not express GRK5 (Violin et al. 2006)" as we believe that in cell disulfide crosslinking provide the most direct and physiologically relevant validation of the interface. In addition, we have read the cited paper (Violin et al. 2006, http://www.jbc.org/content/281/29/20577.long), which indicates that GRK5 (also GRK2, GRK3, GRK6) are ubiquitously expressed, including in HEK cells, but knockdown GRK5 did not have as much effect as knockdown of GRK2/6 on the ability of β2-anerdergic receptor to recruit arrestin.

**Rebuttal Table 1:Cα distances of crosslinking residue pairs at the interface of Arr2-NTSR1 compared with those when both NTSR1 and Arr2 aligned to rhodopsin (Rho) and visual arrestin (Arr1), respectively, in the Arr1-Rho complex orientation. Crosslinking could not happen when Arr2 and NTSR1 form complex in an assembly same as that of the Arr1-Rho complex.**

| Human NTSR1 residue | | Human Arr2residue | | Cα-Cα distance in the Arr2-NTSR1 complex (Å) | Cα-Cα distance with NTSR1 and Arr2 aligned to the positions of Rho and Arr1, respectively, in the Arr1-Rho complex (Å) |
|---|---|---|---|---|---|
| ICL3 (not resolved in the complex structure) | Q273 | N-domain surface | P14 | | |
| | G274 | | P14 | | |
| | Q275 | | K160 | | |
| | C277 | | P14 | | |
| | C277 | | V81 | | |
| | T278 | | K160 | | |
| | G280 | | P14 | | |
| TM6 | R294 | Finger loop | D67 | 14.4 | **23.7*** |
| | R294 | | L68 | 12.0 | **21.0*** |
| | A297 | | D67 | 12.7 | **25.4*** |
| Turn between TM7 and H8 | V367 | | D67 | 9.4 | **19.7*** |
| | S368 | | D67 | 7.9 | **18.9*** |
| | A369 | | D67 | 9.8 | **21.9*** |
| | A369 | | L68 | 11.4 | **18.7*** |
| | A369 | | V70 | 6.2 | **20.7*** |
| | N370 | | D67 | 10.1 | **20.8*** |
| | N370 | | L68 | 12.2 | **17.8*** |
| ICL1 | L94 | Bottom loop | G286 | 6.8 | **23.2*** |
| | Q95 | | G286 | 5.5 | **20.3*** |
| | S96 | | G286 | 6.1 | **17.8*** |
| C-tail | H406 | N-domain surface | P14 | 10.0 | 21.8** |
| | T407 | | P14 | 10.2 | 19.6** |
| | L408 | | P14 | 8.8 | 21.3** |
| | S409 | | P14 | 11.8 | 19.6** |

***Crosslinking could not happen when Cα distance of a residue pair is so far away.**

**The crosslinking at the tail interface may still be possible because of the high flexibility of the C-tail but they are still less likely compared to the Arr2-NTSR1 orientation because of their shorter Cα distances in the crosslinking residue pairs, which are only about half distances of the Arr1-Rho complex.

**Rebuttal Table 2:Cα distances of crosslinking residue pairs at the interface of Arr2-5HTR1A model based on the Arr2-NTSR1 structure, compared with those when both 5HTR1A and Arr2 aligned to Rhodopsin (Rho) and visual arrestin (Arr1), respectively, in the Arr1-Rho complex orientation. Crosslinking could not happen when Arr2 and 5HTR1A form complex in an assembly same as that of the Arr1-Rho complex.**

| Human 5-HTR1A residues | | Human Arr2 residues | | Cα-Cα distance in the Arr2-5HTR1 complex (Å) | Cα-Cα distance with 5-HTR1A and Arr2 aligned to the positions of Rho and Arr1, respectively, in the Arr1-Rho complex (Å) |
|---|---|---|---|---|---|
| TM6 | R339 | Finger loop | D67 | 13.0 | **23.6*** |
| | K342 | | D67 | 14.0 | **23.9*** |
| | R339 | | L68 | 11.4 | **20.8*** |
| | K342 | | L68 | 13.7 | **21.6*** |
| Turn between TM7 and H8 | N404 | | D67 | 8.8 | **19.1*** |
| | K405 | | D67 | 9.8 | **21.6*** |
| | D406 | | D67 | 9.2 | **19.7*** |
| | N404 | | L68 | 11.1 | **16.6*** |
| | K405 | | L68 | 11.7 | **18.7*** |
| | D406 | | L68 | 6.3 | **16.6*** |
| ICL1 | L67 | Bottom loop | R285 | 12.5 | **23.3*** |
| | L67 | | G286 | 12.9 | **25.0*** |

***Crosslinking could not happen when Cα distance of a residue pair is so far away.**

**Rebuttal Table 3: Cα distances of crosslinking residue pairs at the interface of Arr2-5HTR1B model based on the Arr2-NTSR1 structure, compared with those when both 5HTR1B and Arr2 aligned to rhodopsin (Rho) and visual Arrestin (Arr1), respectively, in the Arr1-rhodopsin complex orientation. Crosslinking could not happen when Arr2 and 5HTR1B form complex in an assembly same as that of the Arr1-Rho complex.**

| Human 5-HTR1B residues | | Human Arr2 residues | | Cα-Cα distance in the Arr2-5HTR1B complex (Å) | Cα-Cα distance with 5-HTR1B and Arr2 aligned to the positions of Rho and Arr1, respectively, in the Arr1-Rho complex residues (Å) |
|---|---|---|---|---|---|
| TM6 | R308 | Finger loop | D67 | 11.2 | **22.2*** |
| | R308 | | L68 | 9.7 | **19.3*** |
| | K311 | | D67 | 11.8 | **21.6*** |
| Turn between TM7 and H8 | N373 | | D67 | 8.8 | **19.3*** |
| | D374 | | D67 | 9.8 | **21.5*** |
| | D375 | | D67 | 9.2 | **19.9*** |
| | N373 | | L68 | 11.1 | **18.8*** |
| | D375 | | L68 | 11.7 | **17.0*** |
| | D374 | | V70 | 6.3 | **20.8*** |
| ICL1 | K79 | Bottom loop | R285 | 9.4 | **21.3*** |
| | K79 | | G286 | 8.8 | **23.4*** |

***Crosslinking could not happen when Cα distance of a residue pair is so far away.**

Rebuttal Figure 1

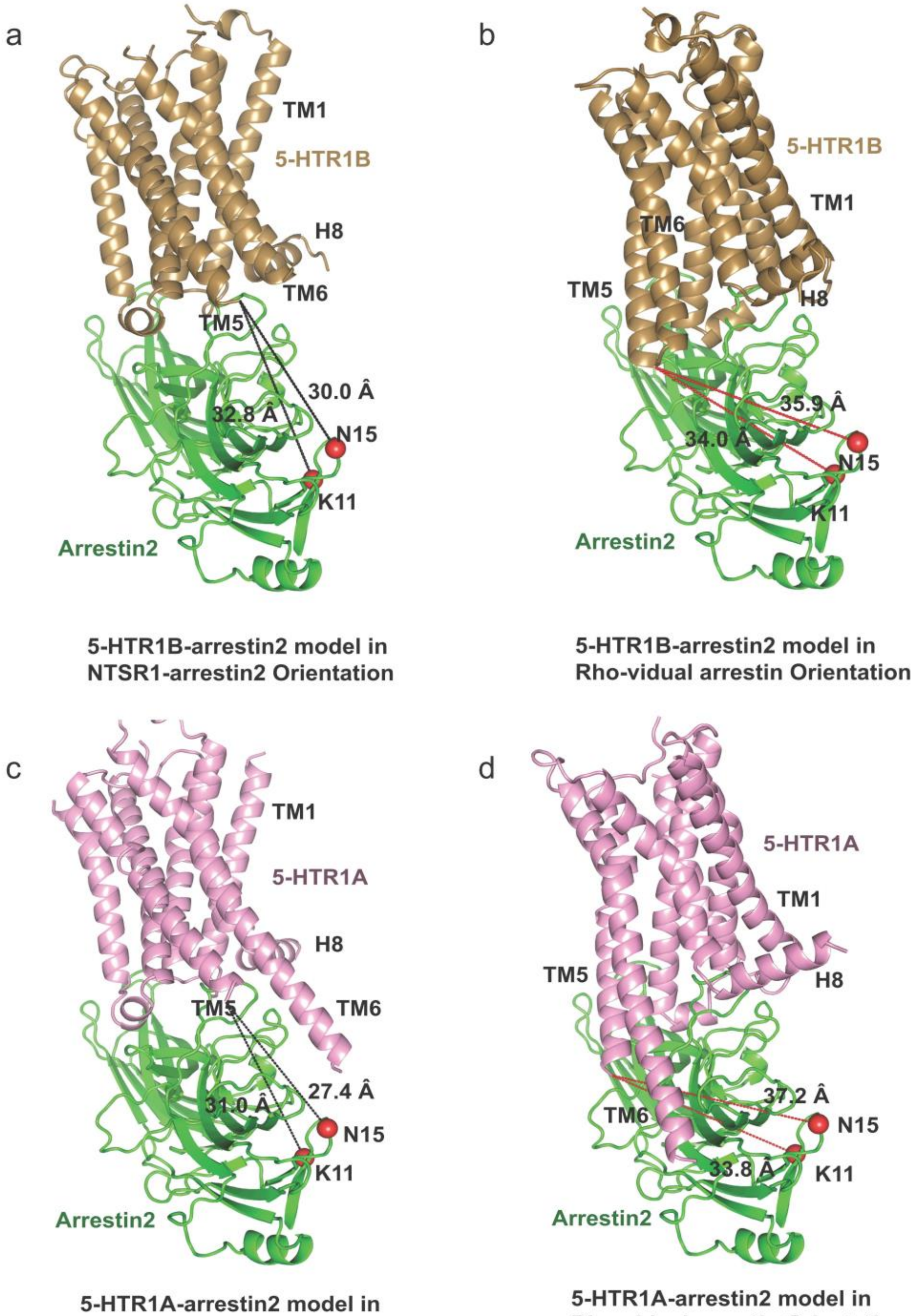


5-HTR1B-arrestin2 model in NTSR1-arrestin2 Orientation

5-HTR1B-arrestin2 model in Rho-vidual arrestin Orientation

5-HTR1A-arrestin2 model in NTSR1-arrestin2 Orientation

5-HTR1A-arrestin2 model in Rho-vidual arrestin Orientation

5) Despite the low resolution of the data, the authors create detailed figures (e.g. Fig. 3a and d) that depict side chains for which configuration is not known, implying a higher resolution than it is. Interpretation of key features like the finger loop are suspect based on the available density. They should try to make figures that do not overextend beyond the quality of the primary data.
>> We agree with the reviewer on this point. As indicated earlier, in the original manuscript, we showed these side chains to indicate the positions of the crosslinking residues but not the actual structure, which we regrettably did not state clearly in the text or in the figure legend. To avoid such confusion, we have re-prepared the relevant figure panels in Figures 3-6. Instead showing the whole residues with side chains, we put little spheres on the backbone trace to show the positions of the specific residues that form disulfide crosslinking with residues at the other side of the interface.

6) There seem to be a lot of typos for a manuscript aimed at this level.
>> We apologize for the typos in the original manuscripts and are grateful for the reviewer to point these out.  We have corrected all those typos and others that we could find in the manuscript.

Editorial issues in order of occurrence:

Abstract: "Arrestins, including visual arrestins and β-arrestins, are critical signal transducers of G-protein-coupled receptors (GPCRs)." Given that arrestin2 or 3 can be knocked out with minor effects, "critical" is probably an overstatement. And the main effect of arresting is probably still receptor desensitization and internalization, not signal propagation. Consider rephrasing.
>> We agree and have modified the text throughout the manuscript.

Abstract: "Sequence analysis and homology modeling suggest that this β-arrestin1–NTSR1 complex structure represents a general paradigm for modeling β-arrestin interactions with a large number of GPCRs." I disagree with this conclusion, as described above. In any event, how do you validate that it is a "large number"?
>>We agree and have rephrased this sentence.

Page 2. "The functions of GPCRs are primarily mediated by G proteins and arrestins, which include two visual arrestins and two β-arrestins." My recommendation would be to adopt the nomenclature "arrestin1-4", which is less confusing and not a misnomer.
>> We agree and have modified the text throughout the manuscript.

Page 2. Typo: "GPCR structure biology"
>>It has been corrected.

Page 3. Typo: "90° different along an axis" change "along" to "around".
>>It has been corrected.

Page 4. Typo: "about four folds" change folds to fold.
>>It has been corrected.

Page 5. "The structure model has been carefully inspected and refined with excellent statistics and geometry (Extended Data Table 1)." It is not possible for a 5 Å structure to have these kinds of statistics, which implies restraints were too high.
>> Structural models with good statistics and geometry can be achieved when Rosetta refinement and manual model rebuilding were iteratively performed (Wang et al. eLife 2016;5:e17219 DOI: 10.7554/eLife.17219). We referenced in the method section that we performed Rosetta refinement and rebuilding but did not mention this in the main text. We have now included this reference in the revised manuscript.

Page 6. "These simulation data suggest that the finger loop as well as the core interface between β-arrestin1 and the receptor is highly dynamic and the assembly of β- arrestin1–NTSR1 complex captured by the cryo-EM structure is probably representing one of many conformational states, which is consistent with the fact that only 5% of total particles were used toward the final cryo-EM structure determination." Given that the map and fit structure is representative of only 5% of the particles, it seems important for readers to understand the extent of heterogeneity in the particle data set. Can the authors analyze eigenvectors, etc. to give a sense of how the various particles in the ensemble differ? Are some arrestins "hanging" instead of interacting with core? Etc.
>> Because of the extreme heterogenicity of the complex preparation, we did not see a clearly defined population of "hanging" particles. I think this is a similar case to the situation of Kobilka/Skiniotis's structure, which uses even a lower percentage of particles (~1%) for their structure determination. In addition, we also analyzed eigenvectors, which indicated the great degree of heterogeneity in the particle data set

Page 6. Typo "side-specific"
>>It has been corrected.

Page 7. "which adopts an extended loop conformation ranging from E66 through L71, " From the map included with the manuscript, I would not be comfortable making this conclusion.
>>We agree and have modified the text.

Page 9. typo "and to a lesser extend"
>>It has been corrected.

Page 9. I strongly disagree with the interpretation of the C-tail density. Phosphorylation at 409 and 410 is not evident. The map shown in Fig. 5b looks quite different the density in the provided EM map, and regardless looks challenging to interpret.
>>We agree and have removed this section from the text in the revised manuscript.

Page 9. "However, the linker region between helix 8 (last residue C383) and Code 1 (S401-T407 or SVSSNHT) contains 17 residues, which is not long enough to cover the distance (about 48 Å based on our structural model) from helix 8 to the site of the tail interface." Again, recommend dispensing with the "Codes". Regardless, the math here seems off because 17 residues in full beta configuration travel about 7 Å between Calpha positions every 2 amino acids, or roughly 60 Å for 17. Thus there is more than

enough distance for "Code 1" to reach the polar core of arrestin.
>>We agree and have removed this section from the text in revised manuscript.

Page 12. The cross linking of the HTR receptors is not very convincing. Examining both rhodospin-arrestin1 complex and the one reported here, a lot of the crosslinks seem compatible with either configuration.
>> This is similar to point #4, which we have addressed earlier.  We agree with the reviewer on that many crosslinking data in ICL3 can be reasonably compatible with both orientations of arrestin in the rhodopsin or NTSR1 complex, including L298C-V81C pairs. However, the crosslinking of ICL3 residue S260C with arrestin residue K11 and N15, as well as the crosslinking of ICL1 residue K79C with arrestin residues R285C or G286C are not compatible with the visual arrestin-rhodopsin orientation, because TM5 is so far away in the back of arrestin N-domain in the rhodopsin orientation, which does not allow S260 to reach over the front side of arrestin for crosslinking. In contrast, these crosslinking data are compatible with the Arr2-NTSR1 orientation, thus supporting our model of coupling Arr2 by ICL3 of 5HT1B (Figure 6).

Page 13. "As β-arrestins are ubiquitously expressed partner proteins for signal transduction of almost all non-visual GPCRs, our structural and biochemical studies of β-arrestin interaction with NTSR1 and 5-HTR1 subfamily members have provided a general paradigm for understanding the mechanistic basis of β-arrestin recruitment and signal transduction by a GPCR." The data in this paper to support this particular hypothesis is weak. Consider revising.
>>We agree and have revised the text.

Page 16. Delete "‘gold-standard’". Everyone writes this about the resolution cutoff, and it implies there is some commission out there that decides what method is gold-standard or not. It also implies that the other EM methods being used are not "gold standard." Shouldn't they all be?
>>We agree and have removed "‘gold-standard’".

Figure Legend 1 panel b. "Symbols and error bars represent mean and s.e.m. of indicated independent numbers of experiments performed in triplicate." Standard deviation should be used instead of SEM for all experiments. The "indicated independent numbers" are not specified. Captions seem terse overall and should ideally include a very brief description of the assay.
>> We obtained data from three independent experiments and think that SEM is correctly used here. Our understanding is that SEM measures how far the sample mean of the data likely differs from a population mean, SD the amount of variability in a dataset.

Figure Legend 1 panel c. Statistics and assay are not described.
>>We agree and it has been added.

Delete Figure 3c.
>>We think that Figure 3C shows conserved negatively charged residues in the finger loop that help to understand the interaction of arrestin with GPCR TMD.

Figure 5c. Just highlight the residues that get phosphorylated and omit the codes. It is unclear what is

going on in the structure. Most consistent with one phosphosite based on the density I see.
>>This part has been removed.

Figure 6. "Our data indicate that the arrestin core structure dynamically shifts". I am not sure what is meant by "dynamically shifts".
>>We have modified the text.

eFigure 1: Delete "gold standard"
>>It has been done.

eFigure 2 typo: "contineous"
>It has been corrected.

efigure 8: "Symbols and error bars represent mean and s.e.m. of indicated independent numbers of experiments performed in triplicate." Standard deviation should be used instead of SEM. The "indicated independent numbers" are not specified.
>> We have performed the experiments three times independently and think that SEM is correctly used here.

Figure 2. NTS density is not observed. Is it reasonable to model it in the figures or coordinates?
>> We can see some density for the backbone trace of NTS and have modified the backbone section of NTS seen in the structure.

Figure 2b. Turn arrows are directionally ambiguous
>>It has been corrected.

Figure 4d. The large number of strong cysteine crosslinks made by K160 and P14 positions indicates to me that there is not a strong sequence preference for this interaction site....a large amount of dynamic behavior.
>>We agree with the comment. K160 and P14 are at the top of the arrestin N-domain, which can crosslink with many residues in ICL3 of the receptor, which is the most dynamics and flexible portion of the receptor.

Make more representative map for Figure 5b.
>>We agree and it has been done.

Delete Figure 5c.
>>We agree and it has been done.

Figure 6e. The strongest crosslink is with Val81, which doesn't seem to argue for placement of the ICL3 loop of HTR receptors in the NTSR configuration as opposed to the more canonical rhodopsin configuration.
>>We agree, and it has been done.

Crosslinking heat maps. Captions could describe what is meant by strong or weak cross linking, or how they were designated.
>>We agree, and it has been done.

**Response to the comments by Reviewer #2:**
Referee #2 (Remarks to the Author):
Review of Yin et al.
„Structure of the Neurotensin Type I Receptor in complex with Arrestin-2“
[Nature manuscript 355343_0]

This is a very interesting manuscript describing the cryo-EM structure of a GPCR together with beta-arrestin-1 (or arrestin-2). The significance of this complex comes from the fact that the two known beta-arrestins interact with and regulate hundreds of GPCRs and in many cases mediate a set of downstream signals that presumably contribute to the important physiological and pharmacological effects that are exerted by these receptors an corresponding drugs. Furthermore, beta-arrestin-mediated signals might in the future be exploited for more specific drugs. Thus, the manuscript does not only represent a conceptual step forward, it also concerns a topic of immense biomedical importance.
>>We thank the reviewer for pointing out that this complex structure is of significant biomedical importance and represents a conceptual step forward in GPCR structural biology.

The authors use the neurotensin receptor 1 (NTSR1) as their model system, which has the advantage that several structures of the receptor alone or with G-protein can be used for comparison. Furthermore, they use their own rhodopsin/arrestin structure as a reference to illustrate similarities as well as differences between these two complexes. This allows them to describe, how flexible these complexes and their interfaces are, explaining how only two beta-arrestins are sufficient to interact with and regulate hundreds of GPCRs. The flexibility of this interaction might furthermore allow the formation of distinct complexes (possibly of distinct functions - but this is not elaborated), which is reflected in the technical difficulty of an apparently large arrays of different complexes that the authors find in their single particle micrographs. Only 5% of the particles could be used in the structural analysis.

In order to enhance the formation of this very flexible complex, the authors combine several approaches: they fuse NTSR1-bArr1 fusion protein structure (GSA-linker), b) NTSR1 carries a N-terminal fusion with BRIL for stabilization, c) beta-arrestin1 is fused C-terminally to the light chain of the so-called Fab30 antibody fragment (which stabilizes the complex formation), d) the Fab30 heavy chain is co-expressed and co-purified so that a full Fab30 is present in the complex, e) they add a positive allosteric modulator, ML314, to stabilize the active state of the receptor, f) they use a pre-activated “3A”-mutant of beta-arrestin1, g) GRK5 is co-expressed to assure phosphorylation of NTSR1 (resulting in different phosphorylation codes – see below).
>>We thank the reviewer for recognizing the highly dynamic and flexible receptor-arrestin interaction, which made the structure determination of a GPCR-arrestin complex a very challenging endeavor. Because of this dynamic nature of the complex, we have extended our works well beyond the structure determination by performing the very extensive set of crosslinking to validate the dynamic assembly of the arrestin-2/NTSR1 complex.

The structure that they obtain shows most of the NTSR1 and of beta-arrestin: NTSR1 R49–T416 (i.e. lacking only the N-terminus the ICL3 (A270 to P292) and the linker region between helix 8 and C-terminal tail, P384 to S404) / β-arrestin1 T6–M352, i.e. lacking the flexible C-terminus. The N-terminally fused BRIL and the small allosteric modulator ML314 are not visible.

A major finding of the manuscript is that compared to the rhodopsin/arrestin structure there is a 90° rotation of beta-arrestin around a vertical axis. Thus, the core interaction between the beta-arrestin central loops and the intracellular pocket of the receptor is maintained, but the "new" orientation of beta-arrestin results in different interactions at the periphery, notably with ICL3 in addition to the phosphorylated receptor C-terminus. Since many GPCRs lack a phosphorylatable C-terminus, but can be phosphorylated in their ICL3, this type of interaction may be of great physiological relevance. This apparent rotational freedom of beta-arrestin may allow it to engage receptors in different modes and thus explain, why just two beta-arrestins are sufficient to interact with hundreds of GPCRs. It probably also explains the technical difficulties in obtaining complexes for structural studies.

As in the rhodopsin/arrestin structure, the beta-arrestin C-edge loop touches the membrane, resulting in an oblique position of beta-arrestin relative to the cell membrane.

To delineate the details of the interaction sites, and because the spatial resolution of the cryEM structure is insufficient to map the side chains, the authors employ extensive C-C crosslinking studies that allow them to nail down contact sites. From these studies, a clear picture emerges that the interaction involves the central loops that interact with the NTSR1 TM core, and a second interaction that involves the NTSR1 C-terminus biding to a positively charged cleft in the beta-arrestin N-terminus. The latter interaction is further studied to understand the phosphorylation code that determines this interaction, with the so-called phosphorylation code 3 (T407-T413) verified by cross-linking.
>>>>We thank the reviewer for recognizing the important results of this manuscript. Our NTSR1-Arr2 structure shows a complex interface different from that observed in the rhodopsin-visual arrestin structure, with a ~90˚ rotation of Arr2 relative to visual arrestin in its rhodopsin complex. This newly found arrestin orientation allows phosphorylated receptor ICL3 to interact with the positively charged arrestin N-domain. The C-lobe of Arr2 is situated to interact with the membrane as does visual arrestin in rhodopsin-visual arrestin complex. We performed in-cell disulfide crosslinking experiments with separately expressed NTSR1 and Arr2, which not only validated the structural model, but also provided additional insights into the dynamic complex interface including the tail interface formed by phosphorylated receptor C-tail or ICL3 with the N-domain of arr2.

The authors then extend their findings to two serotonin receptors, both by modelling and simulations and by cross-linking. These interesting experiments allow them to propose that they have described a generalizable model how GPCRs may interact with beta-arrestins.

Overall, the manuscript is well-written and clear, but some language editing might help to even further improve it.
>>We are grateful for the positive comments on our manuscript and have addressed the language editing issues in the revised manuscript.

I have a few comments and remarks:

– It should be stated more clearly that the structure obtained represents a fusion, and that the two components were quite modified (details see above).

>> We thank the reviewer for this comment. We have now described in both method section and in Introduction how we fused NTSR1-Arr2 complex, and further stabilized the complex, fused Fab30 and Bril at the C-terminus, and N-terminus, respectively, of the complex construct.

– In the introduction, the authors state that the NTSR1 “is an important therapeutic target for a range of diseases including hypotension, obesity, analgesia, drug abuse, cancer, Parkinson’s disease and schizophrenia - they probably want to say that it has been proposed to be such a target.

>> We thank the reviewer to point out that NTSR1 has been proposed to be a target for drug development to treat those diseases and have modified the text in the revised manuscript.

– In the introduction, the authors might want to briefly mention analogies and differences between visual arrestins and non-visual beta-arrestins, such as downstream signaling and a role in internalization. Do they want to suggest that visual arrestin also is involved in signaling and receptor internalization?

>>We thank the reviewer for indicating the analogous and different physiological functions of visual arrestin and beta-arrestin in GPCR signaling. We did not put too many details about the functional differences between those two types of arrestins, but we do mention in Introduction and Conclusions that beta-arrestins (Arr2 and 3) and visual arrestin (arrestin 1) play roles in physiologically different non-visual and visual systems, respectively. The different complex interfaces revealed by our structural studies are consistent with the functional difference between these two types of arrestins. We have modified the text to reflect this point in the revised manuscript.

– In the bindings assays, it is not clear what the authors want to imply with their EC50-values. These are EC50-values for NTS in promoting complex formation, not EC50-values of the interaction itself.

>>The reviewer is correct that these EC50 values are for NTS in promoting complex formation. What we intend to imply is that the preactivated arrestin can increase the binding potency of NTS to the receptor, thus stabilize the complex formation. We have revised the figure legend of Fig 1b accordingly.

– Fig. 3b lacks a negative control - it would be good to see such differences in order to interpret the quantitative aspects of the cross-linking experiments (Ext. Data Fig. 5).

>> We thank the reviewer for this comment. We actually performed disulfide crosslinking for many selected residue pairs at several tentative interface patches. Crosslinking signals are largely different from each residue pairs. Therefore, the residue pairs with no or very weak crosslinking signals serve as negative control for others that show crosslinking signals. To clarify this issue, we have now added a negative control panel in Figure 3b.

– The authors may want to use their structure to propose mechanisms for some intriguing functions of beta-arrestins, such as “action at a distance”/”active conformations after dissociation from receptors” described by Eichel et al. and Nuber et al. They may further want to comment on the possible existence of multiple active conformations of (beta-)arrestins that have been postulated in the rhodopsin field,

but also by Lee&Luttrell, Nuber et al. and others. Given that they use only 5% of the particles for analysis, it would be interesting to know whether any other defined structures might be present in the "rest" and whether these might represent such different structures that have been functionally postulated.
>>We are really interested in various aspects of beta-arrestins' functions and active conformations such as those proposed by Eichel et al. and Number et al. However, due to the multiple functions and therefore the dynamic conformations of arrestins and their GPCR complexes, structural studies are highly challenging. While only 5% of the particles were used, many rejected particles seemed to represent multiple different complex conformations but are very hard to be reconstructed to create decent 3D maps. Thus, at this stage, it is very difficult to speculate on other possible conformational states based on the 3D map reconstruction.

**Response to the comments by Reviewer #3:**
Manuscript: Yin et al., "A complex structure of β-arrestin bound to a G protein-coupled receptor"

Summary: Yin et al. report a reconstruction of beta-arrestin 1 (bARR1) bound to the neurotensin receptor 1 (NTSR1), at a nominal resolution of 4.9Å. The authors generated a stabilized bARR1-NTSR1 complex for structural studies by a multifaceted approach, involving fusion of BRIL to the Nterminus of NTSR1; fusion of bARR1 to the C-terminus of NTSR1 via a 3 amino acid linker; fusion of an antibody fragment to the C-terminus of bARR1; several mutations in both NTSR1 and bARR1 aimed at stabilizing the receptor, including the engineering a NTSR1-bARR1 disulfide bond; phosphorylation of the C-terminal tail of the receptor by GRK5; and incubation of the receptor with both a peptide fragment of neurotensin, and a small molecule agonist, ML314. The authors obtained a cryoEM reconstruction of the resulting complex at 4.9Å, and observed a dramatic rearrangement of bARR1 when compared to the only previous GPCR-bARR complex, of visual arrestin with visual rhodopsin, involving a 90 degree rotation of bARR1 about an axis orthogonal to the membrane. The authors then attempted to validate the interface identified from structural analysis by the introduction of cysteine pair mutations and the identification of resulting disulfide crosslinks. I have been asked to review only the technical aspects of the manuscript relating to structural analysis, which I mention to provide context for the following comments.

**Main impressions:**
The result at the heart of this manuscript is striking, biologically significant, and appears unambiguous at least in the context of the rather extensively engineered construct used for structure determination. I have some concerns regarding how the structural information has been depicted and interpreted, as well as the interpretation of the crosslinking data. These concerns should in my opinion be addressable without further experiments.
>>We are grateful for the positive assessment of the reviewer for the overall quality and significance of our works. This is indeed a very challenging structure which requires specific engineering approaches to stabilize the complex for structural studies. We were pleased to see that we have reached the same conclusions as Kobilka/Skiniotis despite of using different engineering approaches in stabilizing the complex. More importantly, we have validated/extended our structures through extensive in-cell disulfide crosslinking and extended our results further to model ICL3 recruitment of arrestin by tail-less GPCRs. We have coordinated co-submission with Kobilka/Skiniotis and feel that our papers complement

each other and will add tremendous values for readers by publishing these papers together back-to-back.

Our specific responses to each of the reviewer's comments are below.

**Specific comments**:

• NTSR1 is fused to bARR1 with a short 3aa linker from the CT of NTSR1 to the NT of bARR1. In addition, a Fab is fused to the N-terminus of bARR1, and a BRIL module is fused to the NT of the receptor. Given these manipulations, it is essential to validate that the observed arrangement of bARR1 and NTSR1 occurs in a native complex. It is not clear to me that the disulfide crosslinking completely addresses this. Which of the engineered crosslinks would be incompatible with the configuration in the visual arrestin/rhodopsin complex? Many of the mutation pairs which were identified as strongly crosslinking (e.g. N370/L68) here would seem to require substantial conformational change to form a disulfide bond. Given the nominal resolution of the reconstruction this is not surprising, but which of these crosslinks would not be possible in the alternative conformation as observed in the visual arrestin/rhodopsin structure? That is, which can be used to positively confirm that the modeled conformation is what occurs in the native state. It may be worth adding a panel to Ext Data fig 5 to color the weak/medium/strong crosslinking residues on the structure – this would be easier for readers to interpret, in my opinion.

>> We agree with the reviewer's comments and have added a figure to depict the positions of crosslinking residues that are colored based on the weak/medium/strong crosslinking residues on the structure. From this figure, the strongest crosslinking residues are located at the interface. The new Figure (**ED Figure 6b**) is attached in the next page.

• The comparison with the rhodopsin-visual arrestin complex omits one significant difference in the methodology used to obtain the two structures. The rhodopsin-visual arrestin complex was solved using serial femtosecond crystallography with crystals generated by crystallization from the lipidic cubic phase. That is to say, the rhodopsin TMD is embedded in a lipid bilayer, while NTSR1 is in a detergent micelle. Can the authors comment regarding how the interaction with the detergent micelle, as opposed to a true lipid bilayer, might affect the comparison of the two structures? How can the authors be confident that the dramatically different conformation of arrestin observed here is not induced by the presence of the detergent micelle, particularly given arrestin seems to interact with it in a manner that would require conformational change to accommodate the presence of a lipid bilayer?

>> We agree with the reviewer's concern of different conformations in different detergent micelle or in lipid bilayer conditions. Based on the following three points, we would argue that the different orientation of arrestin observed in the complex structure is not induced by different detergent micelle conditions: 1) by superposition of the receptor, we found that the C-edge tip of Arr2 and visual arrestin is at the same height that would interact with the lipid bilayer (Fig 2c); 2) we have validated the new orientation of the Arr2-NTSR1 complex by the extensive in-cell disulfide crosslinking using non-fusion receptor and arrestin. The crosslinking results are consistent with the new orientation of Arr2-NTSR1 complex assembly; 3) through personal communication with Kobilka/ Skiniotis groups, they also found a near 90 degree rotation of Arr2 in the NTSR1 complex, despite using a very different detergent micelle condition, suggest that the different orientation of Arr2 and visual arrestin is not due to the different detergent micelle or lipid conditions.

**ED Figure 6b:** Mapping of the disulfide crosslinking residues at the interface of Arr2-NTSR1 complex. Color code used for the strength of crosslinking signals: strong crosslinking is shown with red (top 75-100%, where 100% mean the strongest crosslinking signal); medium crosslinking, yellow (~50%) ; weak crosslinking, cyan (~25%); and no obvious crosslinking signal, dark blue (0%, no crosslinking signal). The color for the strongest crosslinking was used when a residue was used for multiple crosslinking pairs.

**Extended Data Figure 6b**

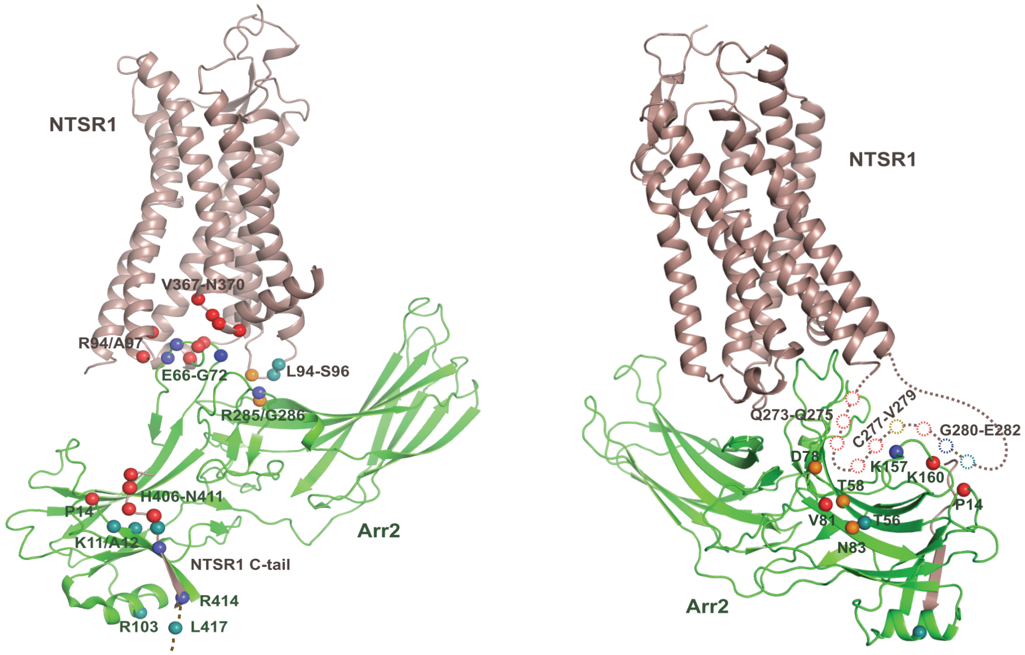


• The authors claim that the resolution of their reconstruction is better than 4.5 Å in the core (presumably this means the transmembrane region). This is not consistent with the visual appearance of the provided density map, which shows no evidence for periodicity of helices, or separation of beta strands. The density for the transmembrane helices in particular appears quite anisotropic. Please provide a 3D-FSC curve to analyze anisotropy, as well as an orientation distribution plot to depict orientation spread. In addition, I would suggest avoid quoting local resolution values determined by ResMap. While these values are useful to qualitatively compare how resolution varies across the structure (which regions are better/worse than average), they are not directly comparable to GS-FSC values used elsewhere. Also, please show the mask used for GSFSC calculation in E.D. Fig 1 C.

>> We agree with the reviewer's comments about the resolution values determined by ResMap. As such, we have removed the sentence that described the resolution value quoted from the ResMap in the revised manuscript. As for the resolution of the core region in our reconstruction, which is not referred to the transmembrane region but a part of the arrestin, which shows a beta-sheet with partially separated strands (see Extended Data Fig. 3 in the revised manuscript). In addition, the GS-FSC analysis of arrestin provided further support for our claim that the highest resolution in our reconstruction should reach 4.5 Å (Extended Data Fig. 2 in the revised manuscript).

By following the reviewer's suggestion, we performed a 3D-FSC and orientation distribution analysis, and we found our previous reconstruction indeed has the anisotropic problem, which resulted from that only a relatively small fraction of top view particles was included in our final reconstruction. To overcome this problem, we added more data from 2D classifications, which contain more top view particles, and carried out an additional round of 3D-classification and auto-refinement. The new reconstruction of the beta-arrestin complex shows a similar overall resolution (4.8 Å) as the previous one but with improved map quality, and the issue of directional resolution anisotropy also gets alleviated (please refer to the updated Extended Figure Fig. 3). The 3D-GSFSC with the used mask have been shown in the revised manuscript.

• In all figures including cryoEM density, please state the threshold used for contouring and other parameters required to reproduce the figure. In particular, if the "carve" option in Pymol has been used, please indicate as much, and give the radius used to carve the map. It appears this has been used quite extensively in Ext Data Fig 2 & Fig 5b, and gives in my opinion a somewhat misleading impression of map quality compared to direct inspection of the map. To be clear, I don't mind if this option is used – just please state clearly any manipulations that change the appearance of the density in the figure legend.
>> All figures related to the density map have been updated with the new density map and all the parameters required to reproduce the figures have been indicated in the revised manuscript. The map figures were prepared with Chimera.

• Please provide more details regarding the 3D classifications – what was the value of the regularization parameter tau? How many classes were used?
>> We used a tau value of 4 and 4-10 classes for the 3D classification. We have added this information into our revised manuscript.

• What program was used for automated particle picking? Relion_autopick?
>> Relion_autopick program was used to pick particles.

• It is indicated that Motioncorr2 was used for frame alignment. This appears to be a typographical error. Please change to MotionCor2.
>>The typo has been corrected.

• Does the conformational spread identified by 3D classification resemble that observed in the MD simulations? Was any rotation of the beta arrestin observed? If not,In Ext fig 1d, it seems like another major (25%) population exhibiting secondary structural features was discarded – does this population have a different global conformation from the one chosen for further analysis?

>>The conformational spread identified by 3D classification (Extended Data Figure 3) indeed resembles that in the MD simulations (Extended Data Figure 5a). Based on the eigenvector analysis, beta-arrestin can rotate differently around the axis vertical to the membrane, even with the same class of 3D classification.

• How was the electrostatic potential map in Fig 5a calculated? I cannot locate it in the methods.
>>This part has been removed from the text as it is associated with the section on the phosphorylation tail, which has been deleted in the revised manuscript.

• The density map has been used to assign the phosphorylation status of individual residues in the C-terminal tail. Inspection of the density suggests the map lacks sufficient features for such a claim.
>>We agree and have removed the figures and text related to the phosphorylation of the receptor.

**We want to thank the three reviewers again for their constructive comments, which have helped us greatly improve the quality of our paper. During the revision of the manuscript, we went back to re-process the EM data, which has improved resolution and map quality. We also rebuilt the structure model, re-prepared the figures, and re-wrote the manuscript to address every comment by the reviewers. The technical merits and validity of our paper have also been improved as the result of our efforts above, for which we hope that the reviewers would agree that our revised manuscript is of high quality for publication as back-to-back papers from the Kobilka/Skiniotis groups.**
